# Imbalance of flight-freeze responses and their cellular correlates in the *Nlgn3^-/y^* rat model of autism

**DOI:** 10.1101/2020.08.27.267880

**Authors:** Natasha J Anstey, Vijayakumar Kapgal, Shashank Tiwari, Thomas C Watson, Anna KH Toft, Owen R Dando, Felicity H Inkpen, Paul S Baxter, Zrinko Kozić, Adam D Jackson, Xin He, Mohammad Sarfaraz Nawaz, Aiman Kayenaat, Aditi Bhattacharya, David JA Wyllie, Sumantra Chattarji, Emma R Wood, Oliver Hardt, Peter C Kind

**Affiliations:** Centre for Discovery Brain Sciences, University of Edinburgh, Edinburgh, EH8 9XD, UK; Simons Initiative for the Developing Brain, University of Edinburgh, Edinburgh, EH8 9XD, UK; Patrick Wild Centre for Autism Research, University of Edinburgh, Edinburgh, EH8 9XD, UK; Dementia Research Institute, University of Edinburgh, Edinburgh, EH8 9XD, UK; Centre for Brain Development and Repair, inStem, National Centre for Biological Sciences, Bangalore, Karnataka, 560065, India; Department of Psychology, McGill University, Montreal, Quebec, H3A 1B1. Canada; The University of Transdisciplinary Health Sciences and Technology, Bangalore, Karnataka, 560065, India

**Keywords:** Fear, freezing, flight, autism, intellectual disability, periaqueductal grey, neuroligin-3

## Abstract

Mutations in the postsynaptic transmembrane protein neuroligin-3 are highly correlative with autism spectrum disorders (ASDs) and intellectual disabilities (IDs). Fear learning is well studied in models of these disorders, however differences in fear response behaviours are often overlooked. Whilst examining fear in a rat model of ASD/ID lacking *Nlgn3*, we observed that they display a greater propensity to exhibit flight responses in contrast to classic freezing seen in wildtypes during fearful situations. Consequently, we examined the physiological underpinnings of this in neurons of the periaqueductal grey (PAG), a midbrain area involved in flight-or-freeze responses. In *ex vivo* slices from *Nlgn3*^-/y^, rats, dorsal PAG (dPAG) neurons showed intrinsic hyperexcitability. Further analysis of this revealed lower magnitude *in vivo* dPAG stimulation evoked flight behaviour in *Nlgn3*^-/y^, rats, indicating the functional impact of the increased cellular excitability. This study provides new insight into potential pathophysiologies leading to emotional disorders in individuals with ASD.

## Introduction

Autism spectrum disorders (ASDs) and intellectual disabilities (IDs) are a complex, heterogeneous group of disorders that are poorly understood in terms of their underlying cellular and circuit pathophysiology. Single-gene mutations account for a large proportion of cases where individuals present with ASD and co-occurring moderate to severe ID (SFARI, McRae *et al.*, 2017) and of these, mutations in synaptic proteins have been repeatedly implicated (Yuen *et al.*, 2017). Mutations in the gene encoding the synaptic protein neuroligin-3 (NLGN3) were originally linked to ASD by Jamain *et al.* (2003) and point mutations in *Nlgn3* have since been shown to be associated with ASD/ID in several studies (Ylisaukko-oja *et al.*, 2005; Talebizadeh *et al.*, 2006; Yu *et al.*, 2011, 2013; Levy *et al.*, 2011; Yanagi *et al.*, 2012; Steinberg *et al.*, 2012; Volaki *et al.*, 2013; Iossifov *et al.*, 2014; Kenny *et al.*, 2014; Mikhailov *et al.*, 2014; Redin *et al.*, 2014; Xu *et al.*, 2014; Yuen *et al.*, 2017; Quartier *et al.*, 2019). The majority of *Nlgn3* mutations identified in humans result in complete or near-complete loss of the NLGN3 protein (Chih *et al.*, 2004; Comoletti *et al.*, 2004; Talebizadeh *et al.*, 2006; Tabuchi *et al.*, 2007; Kenny *et al.*, 2014; Redin *et al.*, 2014; Quartier *et al.*, 2019). NLGN3 is a scaffolding protein expressed at both excitatory and inhibitory synapses where it plays a key role in synaptic development, function and maintenance (Varoqueaux *et al.*, 2006; Budreck and Scheiffele, 2007). Mouse models of both null and disease-causing point mutations in *Nlgn3* lead to behavioural phenotypes as well as alteration synaptic function and plasticity, although the precise nature of these phenotypes differ in a mutation-specific manner (Tabuchi *et al.*, 2007; Etherton *et al.*, 2011; Földy *et al.*, 2013; Zhang *et al.*, 2017; Norris *et al.*, 2019). These synaptic deficits have been shown to underlie circuit and behavioural dysfunction (Tabuchi *et al.*, 2007; Etherton *et al.*, 2011; Baudouin *et al.*, 2012; Rothwell *et al.*, 2014; Polepalli *et al.*, 2017; Hosie *et al.*, 2018). More recently, an *in vivo* study in *Nlgn3* KO mice demonstrated an increase in excitability of neurons in CA2 linked to social cognition deficits (Modi *et al.*, 2019), raising the intriguing possibility that mutations in *Nlgn3* could alter the intrinsic physiology of neurons.

One of the most debilitating aspects of ASD/ID is anxiety and altered emotional responses, however, relatively little is known about the role of NLGN3 in the circuits responsible for fear and emotional learning. Emotional responses have been modelled in animals using fear conditioning paradigms and rat models of ASD and ID have been reported to show deficits in fear learning or extinction (Kim *et al.*, 2008, Lin *et al.*, 2013, Banerjee *et al.*, 2014, Hamilton *et al.*, 2014). Indeed, decreased freezing behaviour during fear conditioning has been shown in *Nlgn3^-/y^* (Radyushkin *et al.*, 2009) mice, but not in *Nlgn3* R451C mice (Chadman *et al.*, 2008; Jaramillo *et al.*, 2017), and results from the rat model are unclear (Hamilton *et al.*, 2014). All of these studies focus on freezing behaviour as the primary readout of fear learning. However, freezing behaviour is not the only fear response exhibited by rodents, or indeed in humans, and hence altered fear expression could be an equally plausible explanation for the reduced freezing. Fight-flight-freeze responses are relatively well characterised, and the decision of which of these responses manifests depends on the context of the fearful situation in which it occurs (De Franceschi *et al.*, 2016).

The importance of the periaqueductal grey (PAG) in regulating the execution of fear responses has been well demonstrated, both by seminal studies from the 1980s and 1990s (Bandler and Carrive, 1988, Bandler and Depaulis, 1988, Tomaz *et al.*, 1988, Schenberg *et al.*, 1990, Zhang, Bandler and Carrive, 1990, Fanselow, 1991, Fanselow *et al.*, 1995) and by more recent work investigating fear circuitry (eg. Johansen *et al.*, 2010; Koutsikou *et al.*, 2015; Wang *et al.*, 2015; Deng *et al.*, 2016; Tovote *et al.*, 2016; Watson *et al.*,, 2016; Evans *et al.*, 2018). Indeed, intra-PAG circuitry allows integration of these inputs, resulting in the expression of active or passive fear responses (Tovote *et al.*, 2016). The role of the PAG in contributing to the pathophysiology of ASD has not yet been elucidated.

This study demonstrates an imbalance in fear responses in *Nlgn3^-/y^*, rats that arises from altered cellular excitability in the PAG.

## Results

### Confirmation of the *Nlgn3^-/y^* rat model

The Nlgn3^-/y^ rats described here contain a zinc-finger nuclease targeted 58 bp deletion in exon 5 (extending into the following intron) of Nlgn3 (**Figure 1A**), and was predicted to cause complete loss of the NLGN3 protein. However, RNA sequencing of WT and Nlgn3^-/y^ cortical tissue revealed only a ~25% loss of Nlgn3 mRNA in Nlgn3^-/y^ rats (**Figure 1C**), and the possible presence of truncated and near-full length isoforms of Nlgn3. The truncated isoform, caused by transcription through the deletion site until a stop codon is reached in the following intron, leads to a potential protein product of 289 amino acids. In addition, a cryptic splice site upstream of the deletion was revealed, with ~25% of RNA-seq reads splicing at this locus to the next exon (**Figure 1D**) leading to a potential protein product 17 amino acids shorter than the full-length protein (**Figure 1B**). As both of these abnormal potential Nlgn3 isoforms are predicted to contain the N-terminus, but not the C-terminus, of NLGN3, we utilised Western blotting to probe for these (**Figure 1E**). We found no presence of NLGN3 protein in Nlgn3^-/y^ cortical homogenate using either C-terminus or N-terminus-specific NLGN3 antibodies (**Figure 1F**), suggesting this abnormal mRNA is not translated to a protein product, and confirming the Nlgn3^-/y^ rat model utilised in this study has complete loss of NLGN3.

**Figure 1.**
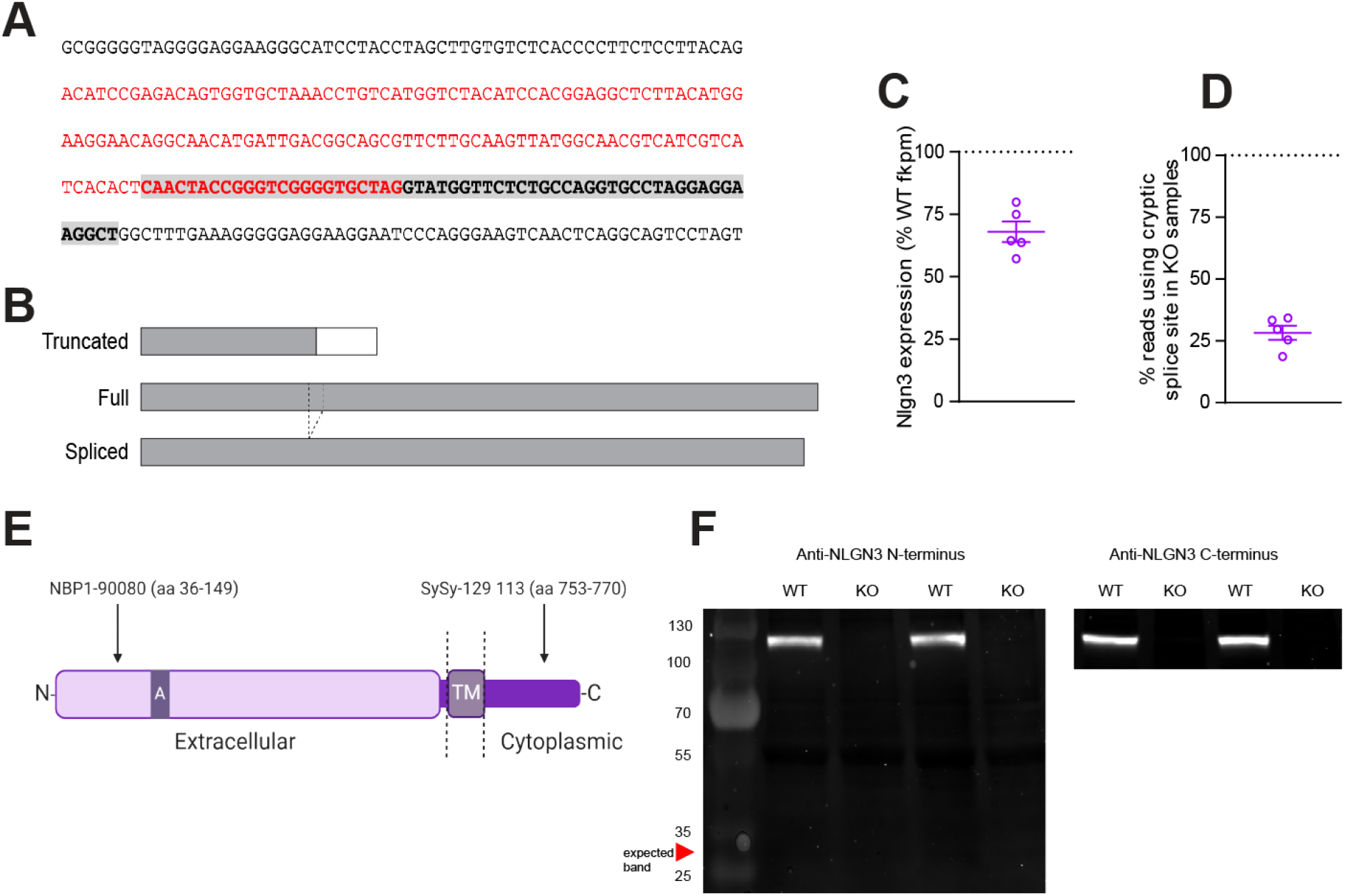
Confirmation of the *Nlgn3^-/y^* rat model. (A) Base sequence of deletion in exon 5. Red text denotes exon, highlighted grey text denotes deletion location. (B) Schematic of potential truncated, full and spliced variants of the NLGN3 protein. Grey regions indicated amino acid sequence shared with the full-length isoform, dotted lines indicate a 17 amino acid section of the full-length *Nlgn3* that is missing in the spliced form. (C) *Nlgn3* mRNA expression in *Nlgn3^-/y^* animals expressed as a percentage of WT (WT n = 6, KO n = 5). (D) Percentage of *Nlgn3^-/y^* RNA-seq reads at the cryptic splice site which splice. (WT n = 6, KO n = 5). (E) Schematic illustrating antibody binding sites to WT NLGN3 protein. (F) Western blot of cortical WT and *Nlgn3^-/y^* tissue using anti-NLGN3 N-terminus (NBP1-90080) and anti-NLGN3 C-terminus (SySy-129 113). No NLGN3 protein of any form was found in *Nlgn3^-/y^* rats (WT n = 2, KO n = 2). Data represented as mean ± SEM, dots represent individual animals.

### *Nlgn3^-/y^* rats exhibit reduced classic freezing behaviour during conditioning and recall phases of auditory fear conditioning

*Nlgn3^-/y^*, mice show reduced freezing during both cued and contextual fear conditioning tasks (Radyushkin *et al.*, 2009). To investigate if this phenotype is also seen in *Nlgn3^-/y^* rats, we trained rats in contextual or auditory fear conditioning paradigms in which they receive three pairings of conditioned (context or tone) and unconditioned (0.9 mA foot shocks) stimuli. In agreement with this data from *Nlgn3^-/y^* mice (Radyushkin *et al.*, 2009), we found that *Nlgn3^-/y^*, rats exhibit reduced freezing behaviour during the conditioning (p < 0.0001, > F_(1, 22)_ = 6.61), recall (p = 0.001, F_(1, 22)_ = 13.36), and extinction (p = 0.0009, F_(1, 22)_ = 14.61) phases of this auditory fear learning paradigm (**Figure 2B-D**). *Nlgn3^-/y^*, rats also exhibited reduced freezing during contextual fear conditioning and recall (**Supplemental Figure 1**).

**Figure 2.**
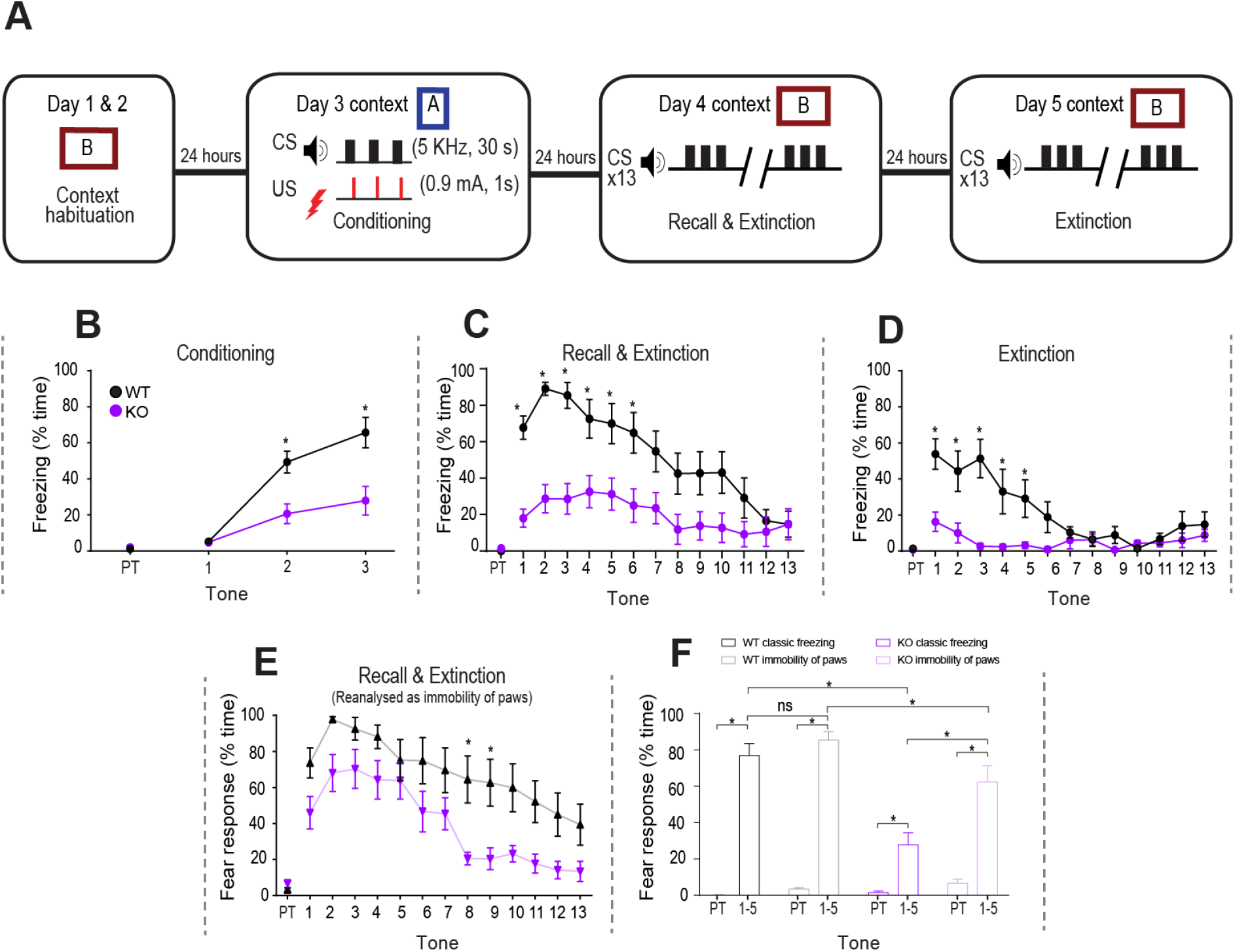
*Nlgn3^-/y^* rats display reduced classic freezing behaviour in an auditory fear conditioning paradigm. (A) Schematic of the auditory fear conditioning protocol. (B) *Nlgn3^-/y^* rats show less classic freezing behaviours during the conditioning phase (p < 0.0001, F_(1, 22)_ = 6.61, repeated measures two-way ANOVA, WT n = 12, KO n = 12). PT: Pre-tone. (C) *Nlgn3^-/y^* rats show less classic freezing behaviours during the recall and extinction phase (p = 0.001, F_(1, 22)_ = 13.36, post-hoc two-way ANOVA, WT n = 12, KO n = 12). (D) *Nlgn3^-/y^* rats show reduced classic freezing behaviours during the second extinction phase (p = 0.0009, F_(1, 22)_ = 14.61, repeated measures two-way ANOVA, WT n = 12, KO n = 12). (E) When analysed as “immobility response” (all four paws unmoving but allowing for movement of head and neck) *Nlgn3^-/y^* rats show significantly increased response to CS in comparison to classic freezing scoring (p = 0.004, F_(1, 22)_ = 13.31, post-hoc two-way ANOVA, KO n = 12). WT rats also show significantly increased paw-immobility response in comparison to classic freezing behaviour (p = 0.019, F_(1, 22)_ = 7.58, post-hoc two-way ANOVA, WT n = 12). Expression of paw immobility response behaviour is significantly lower in *Nlgn3^-/y^* rats in comparison to WT (p < 0.0001, F_(1, 22)_ = 3.26, post-hoc two-way ANOVA, WT n = 12, KO n = 12). (F) Percentage time exhibiting a fear response (defined as either classic freezing (black, purple) or immobility of paws (grey, pink) for pre-tone and average of tones 1-5 of recall shows a significant interaction between genotype, method of scoring and presence of CS (F_(1, 22)_ = 7.52, p = 0.012, three-way ANOVA, WT n = 12, KO n = 12). Both WT and *Nlgn3^-/y^* rats display significant response to the CS (WT classic freezing: p < 0.0001, WT paw immobility: p < 0.0001, KO classic freezing: p = 0.008, KO paw immobility: p < 0.0001, post-hoc Bonferroni-corrected paired t-tests). Scoring method does not affect fear response behaviour during recall for WT rats (p = 0.24, post-hoc paired t-test) however a significantly higher paw immobility response is expressed by *Nlgn3^-/y^* rats in comparison to classic freezing behaviour (p < 0.0001, post-hoc paired t-test). Data represented as mean ± SEM.

Wildtype (WT) cage mates freeze approximately 80% of the time during the first five presentations of the tone during auditory fear conditioning, however, in *Nlgn3^-/y^*, rats this is reduced to around 30% (**Figure 2C**). Similarly, in contextual fear conditioning, WTs freeze approximately 40% of the time, but in *Nlgn3^-/y^*, rats this is reduced to around 10% (**Supplemental Figure 1C**). As freezing behaviour is also reduced during conditioning phases (**Figure 2B**, **Supplemental Figure 1B**) this may be indicative of reduced response to the shock. However, we noticed that the *Nlgn3^-/y^* rats, although not responding with a classic fear response (i.e. no movement except for freezing), were clearly responding to the tone through an immobility of the paws and torso accompanied by jerky head movements, typically in the upward direction. Therefore, we reanalysed the videos for this task, redefining freezing as immobility of the paws and torso but allowing for head movements. This reanalysis revealed that *Nlgn3^-/y^* rats show a fear learning and extinction profile more similar to that of WTs, although still significantly lower (**Figure 2E-F, Supplemental Figure 1D**). Three-way ANOVA analysis between genotype, method of scoring fear behaviour, and CS presentation, revealed a significant interaction between these factors (F_(12, 264)_ = 3.23, p < 0.0001). To explore this interaction further, fear recall (taken to be the mean of the first 5 CS presentations) and pretone were analysed, showing a significant interaction between genotype, method of scoring and presence of CS (F_(1, 22)_ = 7.52, p = 0.012). Additionally, post-hoc paired testing revealed that *Nlgn3^-/y^*, rats display a significantly higher response to the CS when considering immobility of the paws only in comparison to classic freezing (**Figure 2F,** p < 0.0001), however this effect was not seen in WT animals (**Figure 2F**, p = 0.24). In all cases (both genotypes and both methods of scoring), percentage time exhibiting fear response behaviour was significantly higher during the first 5 CS presentations than during the pretone (WT classic freezing: p < 0.0001, KO classic freezing: p = 0.008, WT paw immobility: p < 0.0001, KO paw immobility: p < 0.0001). These findings indicate that *Nlgn3^-/y^* rats, despite showing reduced freezing behaviour, still form the association between tone and shock but may be expressing their fear in a different manner.

### *Nlgn3^-/y^* rats show improved learning of the shocked zone in the active place avoidance task

To further explore a potential role for NLGN3 in fear learning, we employed the active place avoidance (APA) task. The APA task utilises a cylinder with transparent walls and a circular rotating wire grid floor in which rats are required to learn the location of a shock-zone on the grid with respect to stationary cues outside the cylinder (Lesburguères *et al.*, 2016). In this task, one 60° sector of the rotating arena is electrified with a low-ampere shock (0.2 mA), and the rats were given two sessions of 8 trials (10 mins/ trial, **Figure 3A, C**) to learn to position of the shocked zone. When rats initially began training in this task, after receiving one or more low-ampere shocks, 8/9 *Nlgn3^-/y^* rats and 1/9 WT rats tested responded with a jumping behaviour and eventually escaped out of the arena to avoid the shock (**Figure 3B**, p = 0.0034). We therefore had to adapt the arena to include a lid for all subsequent cohorts.

**Figure 3.**
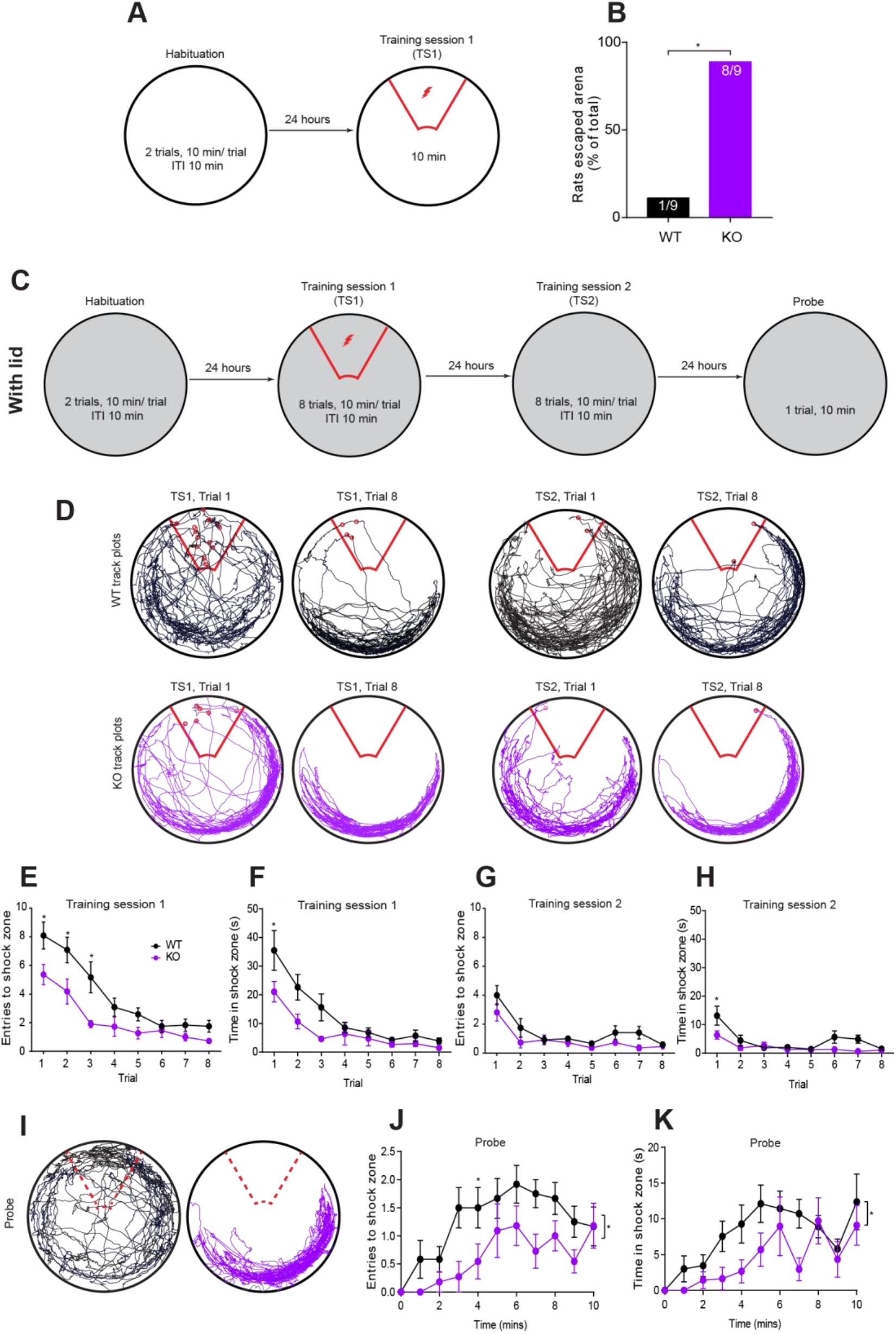
*Nlgn3^-/y^* rats show faster learning and prolonged avoidance of the shock zone in an active place avoidance task. (A) Schematic depicting habituation day and first training session of active place avoidance task (no lid present on arena). (B) 88.9% *Nlgn3^-/y^*, and 11.1% WT rats jumped out of the arena following 0.2 mA foot shocks given during training (p = 0.0034, Fisher’s exact test, WT n = 9, KO n = 9). (C) Schematic of the active place avoidance task, with added lid. (D) Representative track plots for WT and *Nlgn3^-/y^* rats in trials 1 and 8 of training sessions 1 and 2. (E, F) *Nlgn3^-/y^* rats enter the shock zone significantly fewer times during training session 1 (p = 0.0045, F_(1, 21)_ = 10.09, repeated measures two-way ANOVA, WT n = 12, KO n = 11), and spend significantly less time in the shock zone (p = 0.027, F_(1, 21)_ = 5.68 repeated measures two-way ANOVA, WT n = 12, KO n = 11). (G, H) *Nlgn3^-/y^* rats enter the shock zone significantly fewer times during training session 2 (p = 0.044, F_(1, 21)_ = 4.60, repeated measures two-way ANOVA, WT n = 12, KO n = 11), and spend significantly less time in the shock zone (p = 0.025, F_(1, 21)_ = 5.80, repeated measures two-way ANOVA, WT n = 12, KO n = 11). (I) Representative track plots for WT and *Nlgn3^-/y^* rats in the probe trial. (J, K) *Nlgn3^-/y^* rats enter the shock zone significantly fewer times during the probe trial (p = 0.0039, F_(1, 21)_ = 10.51, repeated measures two-way ANOVA, WT n = 12, KO n = 11), and spend significantly less time in the shock zone (p = 0.045, F_(1, 21)_ = 4.53, repeated measures two-way ANOVA, WT n = 12, KO n = 11). Data represented as mean ± SEM.

When tested with a lid on the arena, naïve cohorts of both WT and *Nlgn3^-/y^*, rats learned the location of the shock zone, and could successfully avoid it by remaining in the safe zone by the end of the training sessions (**Figure 3E-H**). As the floor is rotating, this requires active avoidance rather than passively staying out of the shock zone. During habituation to the arena, both WT and *Nlgn3^-/y^* rats showed equal time spent in all sectors of the arena, (**Supplemental Figure 2**) indicating no sections of the arena were inherently aversive before the rats began receiving the low-ampere shocks. *Nlgn3^-/y^* rats displayed enhanced performance in this task; throughout training sessions 1 (TS1) and 2 (TS2) *Nlgn3^-/y^* entered into the shock zone significantly fewer times across trials (**Figure 3**E, G, TS1: p = 0.0045, F_(1, 21)_ = 10.09, TS2: p = 0.044, F_(1, 21)_ = 4.60), and spent significantly less time in this zone (**Figure 3F**, H, TS1: p = 0.027, F_(1, 21)_ = 5.68, TS2: p = 0.025, F_(1, 21)_ = 5.80) in comparison to WT rats.

In the probe trial, no shocks were delivered, allowing us to assess the time taken for the rats to learn that the previously shocked zone is now safe. During the probe trial, *Nlgn3^-/y^* rats displayed significantly prolonged avoidance of previous shock zone. WT rats entered the previous shock zone an average of 13.42 ± 1.55 times, whereas *Nlgn3^-/y^* rats only entered 6.55 ± 1.4 times (**Figure 3J**, p = 0.045, F_(1, 21)_ = 4.53). Additionally, total time spent in previous shock zone was again reduced for *Nlgn3^-/y^*, rats (an average of 4.65 ± 1.12 s in the previous shock zone) in comparison to WTs (an average of 8.23 ± 1.05 s in the previous shock zone) (**Figure 3K**, p = 0.045, F_(1, 21)_ = 4.53). The ability of the *Nlgn3^-/y^*, rats to successfully learn the location of the shock zone agrees with our hypothesis that *Nlgn3^-/y^*, do not have a major learning deficit, indeed, *Nlgn3^-/y^* rats performed better in this task than WT rats. Furthermore, both the exaggerated escape behaviour of *Nlgn3^-/y^* rats seen when this task did not have a lid, along with the increased avoidance during the probe trial, are consistent with altered fear expression in *Nlgn3^-/y^* rats.

### *Nlgn3^-/y^* rats display increased jumping behaviour during a shock-ramp test

One possible explanation for the data described thus far is *Nlgn3^-/y^* rats are hypersensitive to electrical shocks, and this difference in sensitivity leads to atypical fear response behaviour. To test this possibility, we performed a shock-ramp test, in which we gave naïve WT and *Nlgn3^-/y^* rats increasing intensities of foot shocks (0.06 to 1 mA) at fixed intervals and scored and categorised the response behaviour of the rats (**Figure 4A**). Backpedalling and paw withdrawal were the most common initial behaviours elicited when an animal first responded to a foot shock (**Figure 4A**). The minimum shock required to elicit any response was not different between *Nlgn3^-/y^*, and WT rats, an average of 0.072 ± 0.006 mA for WT rats and similarly 0.079 ± 0.01 mA for *Nlgn3^-/y^*, rats (**Figure 4B**, p = 0.13). Furthermore, the minimum shock required to elicit backpedalling behaviour was not different between *Nlgn3^-/y^*, and WT rats; an average of 0.11 ± 0.015 mA for WT rats and 0.087 ± 0.01 mA for *Nlgn3^-/y^*, rats (**Figure 4C**, p = 0.26). This suggests *Nlgn3^-/y^*, rats are not hypersensitive to electrical shocks. However, *Nlgn3^-/y^* rats exhibited significantly more jumping behaviour than WT rats in response to the higher amplitude electrical shocks (**Figure 4D**, p = 0.0081, F_(1, 23)_ = 8.39). These data suggest that *Nlgn3^-/y^*, rats tend to exhibit flight responses to fearful stimuli. At the end of the ramp phase, the shock intensity was decreased to 0.1 mA to test for changes in sensitivity of the animals during the paradigm (**Supplemental Figure 3**). The number of jumps exhibited at 0.1 mA shock intensity were not significantly different on a population level for WT (an average of 0 jumps during the ramp testing and 0.18 ± 0.18 jumps after testing) or *Nlgn3^-/y^*, (an average of 0.07 ± 0.07 jumps during ramp testing and 3.64 ± 2.03 jumps after testing) animals; however, some rats did demonstrate a hypersensitisation to repeated foot shocks (**Supplemental Figure 3**). Nonetheless, these data provide evidence that *Nlgn3^-/y^*, rats do not display altered shock sensitivity yet do show increased flight behaviour (jumping) in response to fearful stimuli.

**Figure 4.**
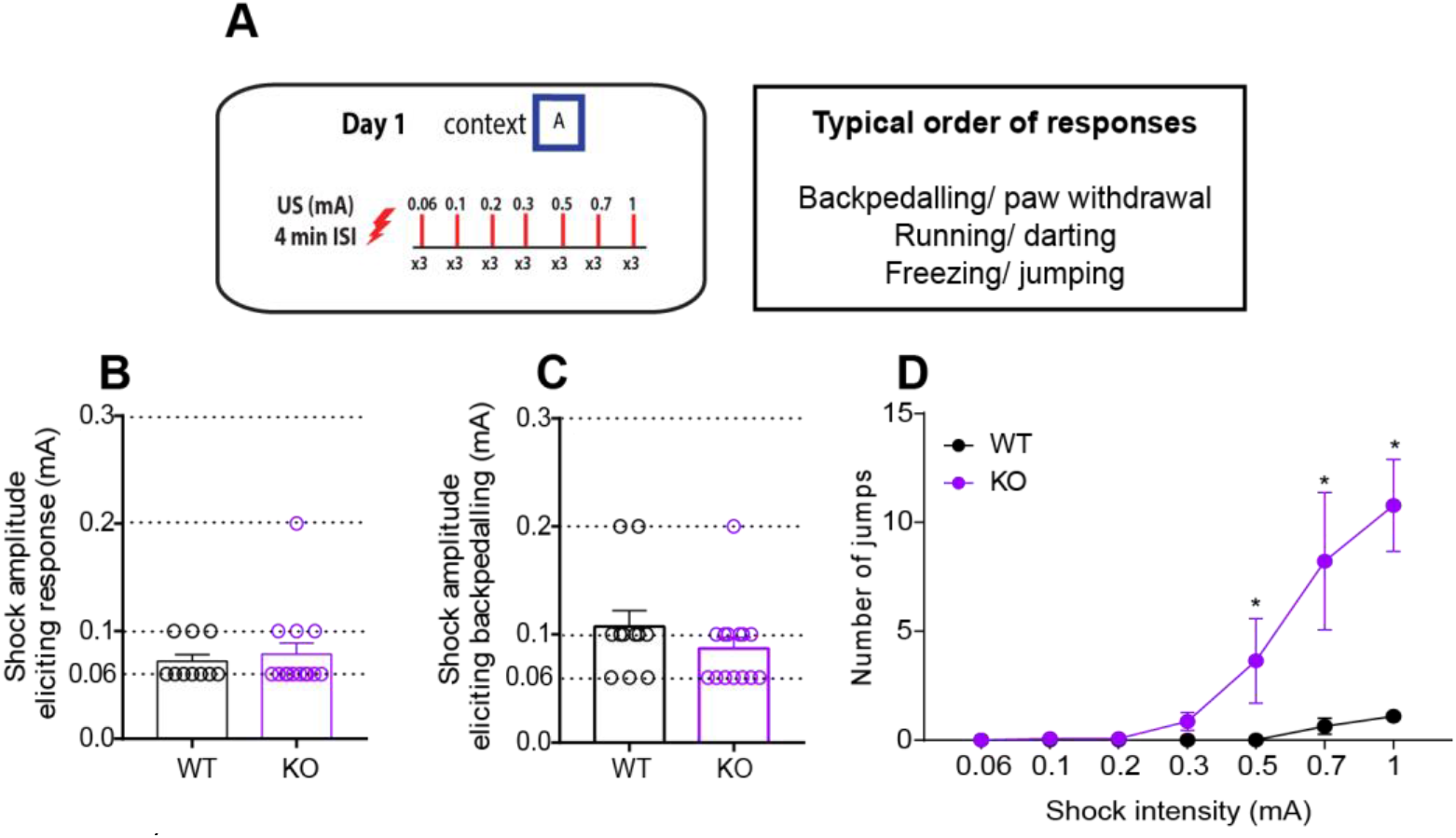
*Nlgn3^-/y^* rats display increased jumping behaviour in response to electrical shocks. (A) Schematic of the shock-ramp test protocol and typical order of responses seen. (B) Lowest shock amplitude required to elicit a response of any kind was not different between WT and *Nlgn3^-/y^* rats (p = 0.13, unpaired t-test, WT n = 11, KO n = 14). (C) Shock amplitude required to elicit backpedalling response was not different between WT and *Nlgn3^-/y^* rats (p = 0.26, unpaired t-test, WT n = 11, KO n = 14). (D) *Nlgn3^-/y^* rats display significantly more jumps in response to increasing intensity electrical foot shocks (p = 0.0081, F_(1, 23)_ = 8.39, repeated measures two-way ANOVA, WT n = 11, KO n = 14).

Data represented as mean ± SEM, dots represent individual animals.

### Dorsal, but not ventral, periaqueductal grey cells in *Nlgn3^-/y^* rats are intrinsically hyperexcitable *ex vivo*

The periaqueductal grey (PAG) is a midbrain region crucial for a host of survival behaviours, including fear responsiveness. Distinct subregions of the PAG are involved in the expression of active or passive fear responses. The dorsal PAG (dPAG) is involved in determining the output of flight responses including jumping and running (Bandler *et al.*, 1985; Bandler and Carrive, 1988; Bandler and Depaulis, 1988; Tomaz *et al.*, 1988; Zhang *et al.*, 1990; Fanselow, 1991; Fanselow *et al.*, 1995; Deng *et al.*, 2016; Assareh *et al.*, 2017). In contrast, the ventral PAG (vPAG) is involved in freezing behaviour (Bandler and Carrive, 1988; Bandler and Depaulis, 1988; Fanselow, 1991; Depaulis *et al.*, 1992; Carrive, 1993; Fanselow *et al.*, 1995; Keay and Bandler, 2001; Deng *et al.*, 2016; Tovote *et al.*, 2016). We hypothesised that the preference for active over passive fear behaviours seen in the *Nlgn3^-/y^*, rats are driven by physiological changes within the PAG. Using whole-cell patch clamp recordings in acute slices from *Nlgn3^-/y^* and WT rats, we measured intrinsic excitability of cells in the dorsal and ventral PAG in response to increasing current steps. We found that cells in the dPAG fired an increased number of action potentials in comparison to WT cells at incremental depolarising current injections (**Figure 5B**, p = 0.018, F_(1, 17)_ = 6.87), and rheobase current was significantly decreased from 68.7 ± 6.5 pA in WT cells to 47.8 ± 5.9 pA in *Nlgn3^-/y^* cells (**Figure 5C**, p = 0.014). There were no changes in the passive membrane properties or action potential threshold (**Supplemental Figure 4**), but the fast-afterhyperpolarisation potential was significantly decreased in dPAG neurons from *Nlgn3^-/y^* rats (**Supplemental Figure 4H**). Conversely, vPAG cells recorded from *Nlgn3^-/y^*, rats fired an equivalent number of action potentials to WT neurons (**Figure 5**F, p = 0.54, F_(1, 17)_ = 0.38) and had an average rheobase current of 44.5 ± 3.9 pA, which was comparable to that of WT rats: 36.6 ± 2.9 pA (**Figure 5G**, p = 0.40). *Nlgn3^-/y^* vPAG neurons did, however, display increased membrane time constants (**Supplemental Figure 4**). The observed hyperexcitability of dPAG cells in *Nlgn3^-/y^* rats may explain the increased flight and decreased freezing behaviour seen in these rats.

**Figure 5.**
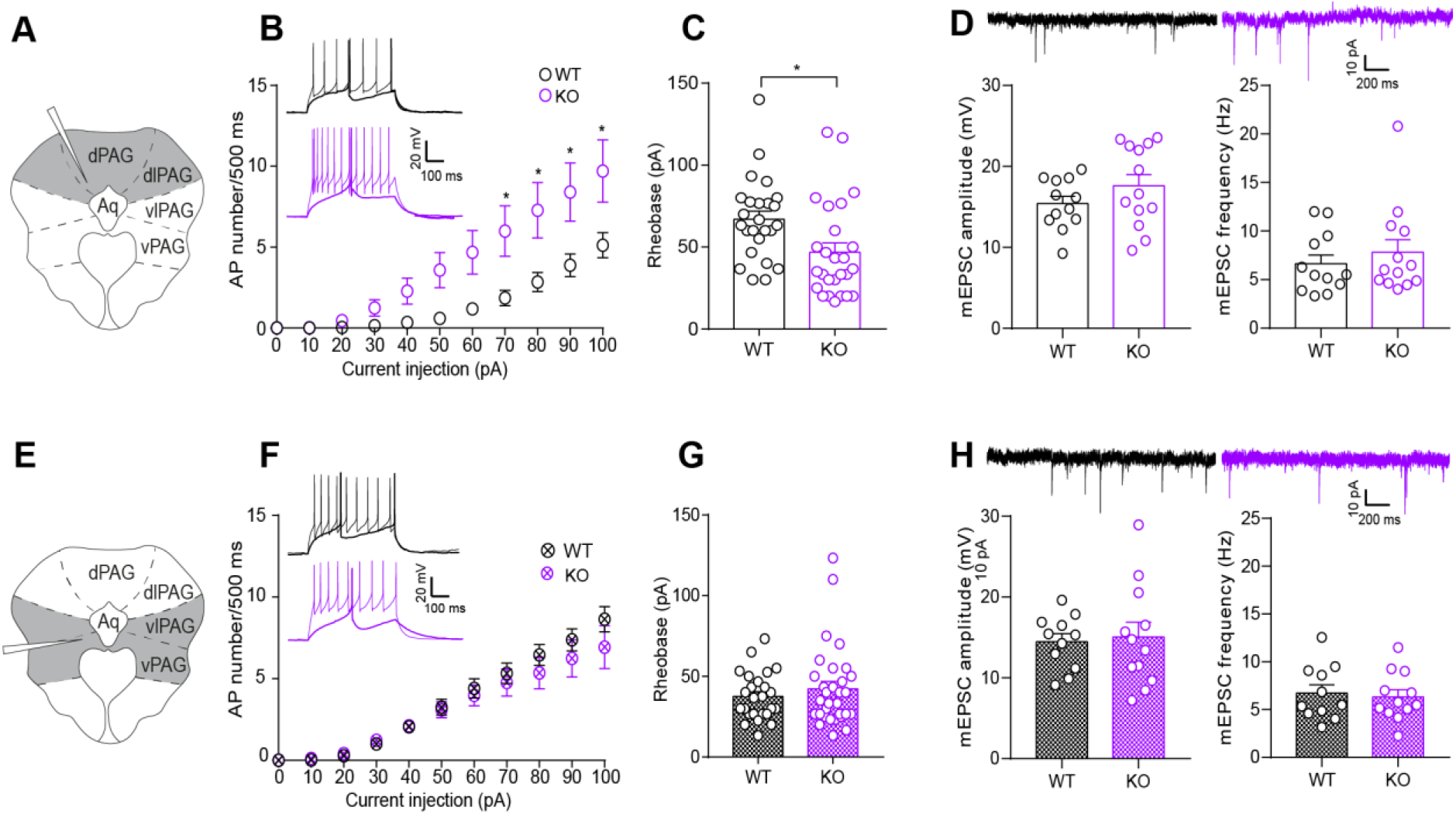
Hyperexcitability of dorsal, but not ventral PAG neurons in *Nlgn3^-/y^* rats. (A, E) Schematics of PAG slice indicating area recorded from in grey. (B) dPAG cells from *Nlgn3^-/y^* rats fire increased numbers of action potentials in response to increasing current injection steps (p = 0.018, F_(1, 17)_ = 6.87, repeated measures two-way ANOVA, WT n= 25 cells/ 10 rats, KO n = 26 cells/ 9 rats). Representative traces of rheobase and +100 pA steps for WT (black) and *Nlgn3^-/y^* (purple) dPAG cells. (C) dPAG cells from *Nlgn3^-/y^* rats have lower rheobase potential than WT (p = 0.014, GLMM, WT n= 25 cells/ 10 rats, KO n = 26 cells/ 9 rats). (D) No change in mEPSC amplitude or frequency of dPAG neurons in *Nlgn3^-/y^* rats compared to WT (amplitude: p = 0.28, frequency p = 0.61, GLMM, WT 12 cells/ 6 rats, KO 13 cells/ 6 rats). Representative traces of mEPSCs of dPAG cells from WT (black) and *Nlgn3^-/y^* (purple) rats. (F) vPAG cells from *Nlgn3^-/y^* and WT rats fire comparable numbers of action potentials in response to increasing current injection steps (p = 0.54, F_(1, 17)_ = 0.38, repeated measures two-way ANOVA, WT n = 24 cells/ 9 rats, KO n = 28 cells/ 10 rats). Representative traces of rheobase and +100 pA steps for WT (black) and *Nlgn3^-/y^* (purple) vPAG cells. (G) vPAG cells from *Nlgn3^-/y^* and WT rats have comparable rheobase potentials (p = 0.40, GLMM, WT n= 24 cells/ 9 rats, KO n = 28 cells/ 10 rats). (H) No change in mEPSC amplitude or frequency in vPAG neurons *Nlgn3^-/y^* rats compared to WT (amplitude: p = 0.78, frequency p = 0.88, GLMM, WT 11 cells/ 5 rats, KO 12 cells/ 6 rats). Representative traces of mEPSCs of vPAG cells from WT (black) and *Nlgn3^-/y^* (purple) rats. Data represented as mean ± SEM, dots represent individual cells.

In addition to intrinsic excitability, the excitability of a neuron depends on the synaptic input it receives. We measured mEPSC amplitude and frequencies in dorsal and ventral PAG cells using whole-cell patch clamp recordings in acute slices from *Nlgn3^-/y^* and WT rats. We found that cells recorded from *Nlgn3^-/y^*, and WT rats had comparable mEPSC amplitudes and frequencies in both dPAG (WT average amplitude/frequency: 15.41 ± 0.91 pA/ 6.64 ± 1.27 Hz, KO average amplitude/frequency: 17.59 ± 1.85 pA/ 7.58 ± 1.54 Hz, **Figure 5D**, amplitude: p = 0.28, frequency: p = 0.61) and vPAG (WT average amplitude/frequency: 13.94 ± 0.91 pA/ 6.91 ± 1.27 Hz, KO average amplitude/frequency: 15.05 ± 1.59 pA/ 6.33 ± 1.01 Hz, **Figure 5H**, amplitude: p = 0.78, frequency: p = 0.88). Together, these data suggest that dPAG cells are intrinsically hyperexcitable, but do not appear to receive altered excitatory synaptic input.

### *Nlgn3^-/y^* rats display normal tone-evoked LFP amplitudes in the PAG during fear recall

Reduced freezing behaviour during fear recall and extinction has been shown to be correlated with reduced tone-evoked local field potential (LFP) amplitudes (event-related potentials, ERPs) in the PAG (Watson *et al.*, 2016). Additionally, synaptic activity of individual neurons has been linked to the onset of both spontaneous and evoked LFPs *in vivo* (Haider *et al.*, 2016). In order to determine if this was true for the decreased freezing behaviour seen in *Nlgn3^-/y^* rats (**Figure 2**, **Supplemental Figure 1**) we unilaterally implanted naïve WT and *Nlgn3^-/y^* rats with electrodes in the dPAG (see Methods), and recorded LFPs evoked by the CS tone during auditory fear recall (**Figure 6**). We observed again that *Nlgn3^-/y^*, rats display less freezing behaviour in comparison to WTs across all 13 tones of fear recall and extinction, (**Figure 6E**, p = 0.032, F_(1, 13)_ = 1.96), and instead exhibited limb immobility with head movements as described in **Figure 2** (**Figure 6F**). Three-way ANOVA analysis revealed a significant interaction between genotype and method of scoring fear behaviour (p = 0.014, F_(1, 13)_ = 8.04), and post-hoc two-way ANOVAs indicated no difference in the percentage of time WT animals spend exhibiting classic freezing in comparison to with their paws immobile (p = 0.12, F_(1, 6)_ = 3.39). In contrast, as was seen in **Figure 2**, *Nlgn3^-/y^* rats showed a significant difference between the expression of these two behaviours (p = 0.001, F_(1, 7)_ = 29.75).

**Figure 6.**
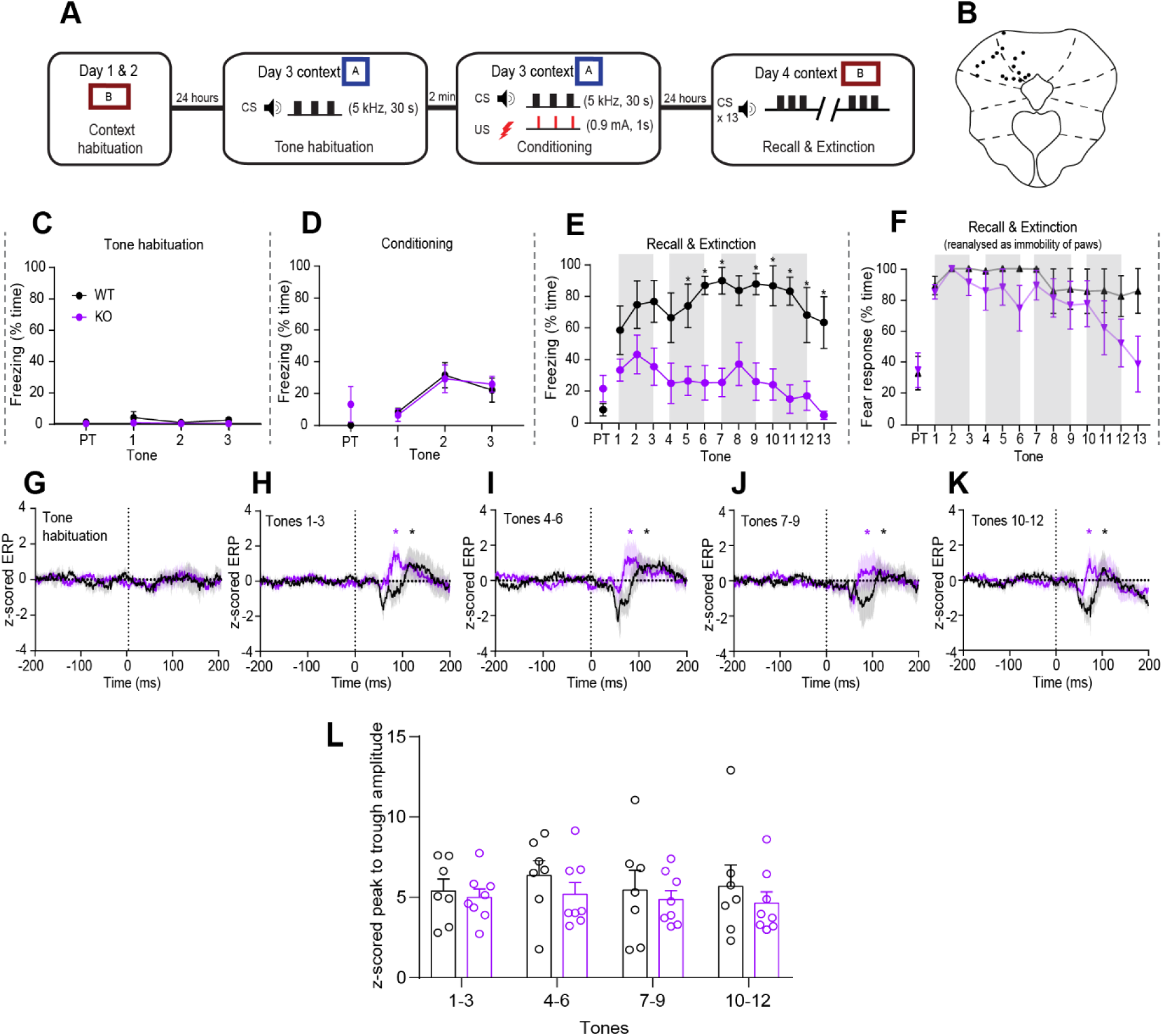
*Nlgn3^-/y^* rats show similar dPAG ERP amplitudes to WT rats during an auditory fear conditioning paradigm. (A) Schematic of auditory fear conditioning paradigm including tone habituation session. (B) Approximate locations of recording sites in the PAG. Black dots represent lesion site after electrode removal for an individual animal. (C) Almost no classic freezing behaviour was seen during the tone habituation session (p = 0.13, F_(1, 13)_ = 2.63, repeated measures two-way ANOVA, WT n = 7, KO n = 8). PT: Pre-tone. (D) WT and *Nlgn3^-/y^* rats display similar levels of freezing behaviour during auditory fear conditioning (p = 0.54, F_(1, 13)_ = 0.74, repeated measures two-way ANOVA, WT n = 7, KO n = 8). (E) *Nlgn3^-/y^* rats display significantly lower freezing behaviour during fear recall in comparison to WT cage-mates (p = 0.0015, F_(1, 13)_ = 16.12, repeated measures two-way ANOVA, WT n = 7, KO n = 8). Grey boxes represent tone-responses that have been averaged for panels H-K. (F) When analysed as “immobility response” (all four paws unmoving but allowing for movement of head and neck) *Nlgn3^-/y^* rats show no difference in this behaviour during fear recall in comparison to WT controls (p = 0.014, F_(1, 13)_ = 8.036 (analysis method x genotype), p = 0.28, F_(12, 156)_ = 1.21 (tone x analysis method), p = 0.093, F_(12, 156)_ = 1.62 (tone x genotype), three-way ANOVA, WT n = 7, KO n = 8). (G) No significant ERPs during tone habituation for WT (n = 7, p = 0.25, paired t-test) or *Nlgn3^-/y^* (n = 8, p = 0.093, paired t-test) rats. Data represented as mean (solid line) ± SEM (translucent shading). (H-K) Significant ERPs were observed in both WT and *Nlgn3^-/y^* z-scored average ERP waveforms after CS onset for tones 1-3 (WT: p = 0.0032, KO: p = 0.0099), 4-6 (WT: p = 0.0030, KO: p = 0.0084), 7-9 (WT: p = 0.029, KO: p = 0.004) and 10-12 (WT: p = 0.0158, KO: p = 0.0046, paired t-tests, WT n = 7, KO n = 8). Data represented as mean (solid line) ± SEM (translucent shading). Dotted lines represent tone onset. (L) No significant difference in z-scored ERP peak to trough amplitude during fear recall in WT and *Nlgn3^-/y^*, rats (p = 0.42, F_(1, 13)_ = 0.73, repeated measures two-way ANOVA, WT n = 7, KO n = 8). Data represented as mean ± SEM, dots represent individual animals.

Due to previously identified correlation between freezing behaviour during fear extinction and PAG ERP amplitude (Watson *et al.*, 2016), we hypothesised CS-evoked ERPs would be reduced in *Nlgn3^-/y^* rats. However, we observed robust ERPs in the PAG of both *Nlgn3^-/y^* and WT rats during fear recall (**Figure 6H-K**), the amplitude of which did not differ between WT and *Nlgn3^-/y^* rats (**Figure 6L**, p = 0.42, F_(1, 13)_ = 0.73), despite significantly different freezing responses exhibited by the two genotypes (**Figure 6E**). It was noted, however, that the peak to trough duration of these ERPs was significantly shorter in *Nlgn3^-/y^*, rats in comparison to WTs (**Supplemental Figure 5**), although the biological relevance of this finding is currently unclear. In a subset of the same rats (WT n = 5, KO n = 7), we recorded LFPs during the tone habituation session to determine if ERPs were triggered by the unconditioned tone. Contrary to the ERPs seen after conditioning, no ERPs were observed in response to tone during tone habituation (**Figure 6G**), and no freezing behaviour was exhibited by either genotype (**Figure 6C**). These results indicate that *Nlgn3^-/y^* rats display robust CS-evoked ERPs in the PAG during fear recall, comparable in amplitude to that of WT rats, despite significant changes to fear response behaviour. These data suggest that CS-evoked ERPs in the PAG reflect overall fear state elicited by the tone and are not indicative of the type of fear response behaviour (i.e. freezing, flight) exhibited.

### *Nlgn3^-/y^* rats show increased jumping behaviour in response to *in vivo* dPAG stimulation

Seminal studies from the 1980s and 1990s have shown dPAG electrical or chemical stimulation evokes robust escape responses such as running and jumping (Bandler and Carrive, 1988, Bandler and Depaulis, 1988, Tomaz *et al.*, 1988, Schenberg *et al.*, 1990, Zhang *et al.*, 1990, Fanselow, 1991, Fanselow *et al.*, 1995). Having seen increased escape behaviour in *Nlgn3^-/y^*, rats in response to increasing electrical shocks (**Figure 4**), and increased intrinsic excitability of dPAG neurons from *Nlgn3*, rats (**Figure 5**), we investigated the effect of direct increasing electrical stimulation of the dPAG in *Nlgn3^-/y^* rats and WT cage-mates. We bilaterally implanted stimulating electrodes in the dPAG of naïve *Nlgn3^-/y^* and WT rats (see Methods, **Figure 7B**), and, after recovery from surgery, stimulated the PAG in increasing intensity steps from 30-75 μA (0.1 ms pulses,100 Hz train, 2 s total duration; see methods, Kim *et al.*, 2013) to observe behavioural responses (**Figure 7A**). We observed that dPAG stimulation resulted in an immediate post-stimulation activity burst lasting 1-5 seconds, followed by freezing (reviewed in Brandão and Lovick, 2019). We observed that a significantly higher percentage of *Nlgn3^-/y^* rats exhibited successful escapes from the arena during the increasing dPAG stimulations (p < 0.0001, **Figure 7C**), and furthermore a significantly higher percentage of *Nlgn3^-/y^* rats exhibited jumping behaviour at dPAG stimulations of 60 μA (20% WT, 77.8% KO), 65 μA (20% WT, 66.7% KO) and 70 μA (20% WT, 77.8% KO) in comparison to WT rats (p = 0.0065, **Figure 7D**). *Nlgn3*, rats also displayed reduced overall classic freezing behaviour throughout the experiment (p = 0.025, F_(1, 12)_ = 6.58, **Figure 7E**). To control for the possibility that non-specific brain stimulation was causing an increase in escape behaviour in the *Nlgn3^-/y^* rats, a subset of animals (2 WT and 2 *Nlgn3^-/y^*) were bilaterally implanted with stimulating electrodes in primary somatosensory cortex. Given incremental stimulation of primary somatosensory cortex, these rats displayed no jumping or escape-like behaviour at any time during the behavioural assessment. Classic freezing behaviour was also scored throughout this protocol, and although some immobility indistinguishable from true fearful freezing was observed (**Supplemental Figure 6**) the same profile of post-stimulation activity burst followed by freezing behaviour was not seen. These data provide evidence that the threshold for PAG-evoked jumping behaviour is reduced in *Nlgn3^-/y^* rats, and supports our hypothesis that increased intrinsic excitability of dPAG cells in *Nlgn3^-/y^*, rats causes a circuit bias to favour the output of flight rather than freezing behaviours.

**Figure 7.**
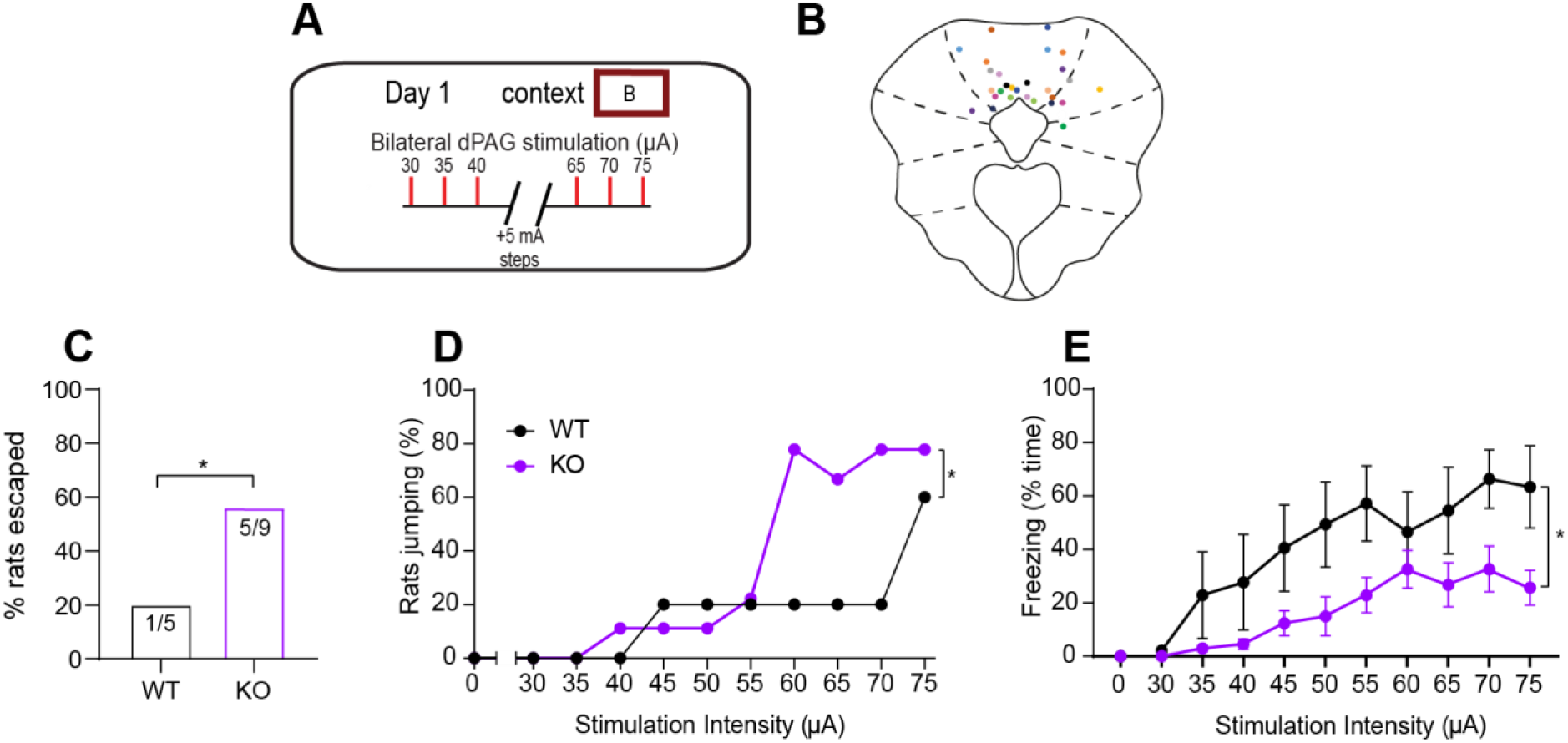
*Nlgn3^-/y^* rats show increased jumping behaviour in response to in vivo dPAG stimulation. (A) Schematic depicting dPAG stimulation protocol. (B) Location of implanted stimulating electrodes. Coloured dots represent lesion sites (bilateral) of individual animals. (C) Significantly more *Nlgn3^-/y^* rats successfully escaped the arena following dPAG stimulation in comparison to WT rats (WT n = 5, KO n = 9, p < 0.0001, Fisher’s exact test). (D) A higher percentage of *Nlgn3^-/y^* in comparison to WT rats display jumping behaviour when increasing bilateral dPAG stimulations (p = 0.0065, Fisher’s exact test, WT n = 5, KO n = 9). Data represented as mean %. (D) Classic freezing behaviour is reduced in *Nlgn3^-/y^* rats (p = 0.025, F_(1, 12)_ = 6.58, repeated measures two-way ANOVA, WT n = 5, KO n = 9). Each data point is mean % time freezing over the entire 3-minute interval following stimulation ± SEM.

## Discussion

In this study we show that the *Nlgn3^-/y^* rat model of ASD/ID has abnormal responses to fearful stimuli in both fear conditioning and as a direct result of foot shocks. *Nlgn3^-/y^*, rats display increased flight and decreased freezing behaviour in response to fearful stimuli, in comparison to WT cage-mates. We also provide evidence that learning and memory are not impaired in *Nlgn3^-/y^*, rats. Furthermore, despite significantly reduced freezing behaviour displayed by *Nlgn3^-/y^*, rats during fear recall, the amplitude of tone-evoked LFPs in the PAG are unaffected. Correspondingly, excitatory synaptic inputs to cells in the PAG of *Nlgn3^-/y^* rats are comparable to those of WTs. We also show that dPAG cells in *Nlgn3^-/y^*, rats have increased intrinsic cellular excitability *ex vivo*, and that *Nlgn3^-/y^*, rats exhibit atypical responses to direct dPAG stimulation *in vivo*. To our knowledge, neither imbalance of flight-freeze responses nor electrophysiological changes in the PAG have been previously reported in any model of ASD or ID.

### Abnormal fear responses in the Nlgn3^-/y^ rat model

Fear conditioning and recall is often used to assess emotional learning in ASD/ID models, using the quantification of freezing behaviour as a proxy for the memory of the CS-US association. We find *Nlgn3^-/y^* rats display less freezing behaviour (defined as no movement except for respiration) during both auditory and contextual fear recall than WT rats. Taken in isolation, these data could be interpreted as reduced fear learning and/or memory in *Nlgn3^-/y^* rats. However, reanalysis of these data revealed that *Nlgn3^-/y^* rats stop exploratory behaviours following onset of the tone, and respond by staying fixed in the same location within space but moving the head and neck. This type of fear behaviour has been reported before in rats confronted with a snake (Uribe-Mario *et al.*, 2012, Calvo *et al.*, 2019). This suggests *Nlgn3^-/y^* rats do form an association between the CS and US, but are expressing their fear differently to WT rats. We did not observe escape behaviour during this task, likely because the arena was fully enclosed with no possible escape route. Radyushkin *et al.* (2009) described reduced freezing in the *Nlgn3^-/y^* mouse, however no further investigation was made into the fear responses of these mice, so it is not known whether these two models of *Nlgn3* deficiency display converging phenotypes. Interestingly, Hosie *et al.* (2018) reported increased jumping behaviour in *Nlgn3* R451C mice during a social interaction task, consistent with our findings.

Further insight into the fear responses and learning of *Nlgn3^-/y^* rats was seen in direct response to electrical foot shocks. Active place avoidance (APA) and a shock-ramp paradigm revealed *Nlgn3^-/y^*, rats exhibit escape behaviours in response to foot shocks much more readily than WT controls. However, *Nlgn3^-/y^* rats were able to efficiently learn the location of a shock-zone in the APA task once escape routes were blocked. Moreover, shock sensitivity testing revealed that *Nlgn3^-/y^* rats are not hypersensitive to electrical shocks, but again show increased jumping (flight) responses. These data further support our hypothesis that *Nlgn3^-/y^*, rats do not display associative learning deficits, but preferentially exhibit flight over freezing behaviour in response to fear.

### Cellular correlates of flight-freeze responses

Control of flight and freeze responses to fear are known to involve the dorsal and ventral PAG. Low intensity electrical stimulation of the dorsal PAG has been shown to elicit freezing responses (Schenberg *et al.*, 1990, Vianna *et al.*, 2001) and higher stimulation to elicit flight responses (Bandler and Carrive, 1988, Bandler and Depaulis, 1988, Tomaz *et al.*, 1988, Schenberg *et al.*, 1990, Zhang *et al.*, 1990, Fanselow, 1991, Fanselow *et al.*, 1995), whereas stimulation of the ventral PAG has been shown to elicit freezing responses rather than flight responses (Zhang *et al.*, 1990, Fanselow, 1991, Fanselow *et al.*, 1995). Correlating with the increased flight behaviour seen in *Nlgn3^-/y^* rats, we observe increased intrinsic cellular excitability in the dorsal, but not ventral, PAG in slices from naïve *Nlgn3^-/y^*, rats. These changes in intrinsic excitability in the dorsal PAG are likely to affect the excitation/inhibition balance within the PAG, bringing the resting state of *Nlgn3^-/y^*, rats closer to the “threshold” of eliciting an escape response (Evans *et al.*, 2018).

The increase in firing frequency in the dPAG of *Nlgn3^-/y^* rats appears to be a result of reduced fast afterhyperpolarisation potential (fAHP). fAHP is mediated by Ca^2+^-activated large-conductance K^+^ channels (BK) which act to hyperpolarise the membrane and reduce neuronal firing (Springer *et al.*, 2015). BK channel open-probabilities have been shown to be decreased in another model of ASD/ID, the *Fmr1^-/y^*, mouse, leading to increased neuronal excitability (Deng and Klyachko, 2016). This presents an interesting future research avenue into the function of BK channels in the *Nlgn3^-/y^* rat.

CS-evoked ERPs recorded from the PAG during fear conditioning and recall have been reported to reduce in amplitude during CS-US extinction, correlating with reduction in freezing behaviour (Watson *et al.*, 2016). However, we observe that despite exhibiting significantly less classic freezing behaviour than WT rats, the PAG ERP amplitudes in *Nlgn3^-/y^*, rats does not differ from that of WTs. This suggests that CS-evoked ERPs in the PAG reflect the presence of fear, but are unrelated to the type of response the rat is exhibiting. The presence of robust CS-evoked ERPs in *Nlgn3^-/y^* rats supports our hypothesis that these rats acquire learned fear of the tone despite the significantly reduced freezing behaviour they exhibit. Several studies have reported a strong positive correlation between synaptic input to a neuron and LFP magnitude (Haider *et al.*, 2016, Wright *et al.*, 2017, Arroyo *et al.*, 2018). Consistent with this, we found that miniature excitatory post synaptic currents (mEPSCs) were not altered in either dorsal or ventral PAG cells recorded *ex vivo* from *Nlgn3^-/y^*, rat slices, suggesting that excitatory synaptic input to these PAG neurons was not altered.

Additionally, we note that the peak-to-trough duration of CS-evoked ERPs in the dPAG of *Nlgn3^-/y^* rats are significantly shorter than those in WTs. The biological relevance of this finding is not clear, however as voltage-gated ion channels have been suggested to affect LFP waveform (Reimann *et al.*, 2013; Ness *et al.*, 2016, 2018), the altered BK channel conductance implicated by the reduced fAHP observed in dPAG neurons *ex vivo* may be contributing towards this phenotype. The broader tone-evoked LFP we observe in the dPAG of *Nlgn3^-/y^*, rats during fear recall may be reflective of longer, more sustained activity in the PAG. Further experimentation is required to understand this.

Finally, we show that *in vivo* dPAG stimulation elicits escape and flight responses in a significantly higher percentage of *Nlgn3^-/y^*, rats than of their WT cage-mates. If intrinsic excitability of dPAG cells in *Nlgn3^-/y^*, rats is increased, additional stimulation of this brain region may cause flight responses to be elicited at a lower threshold than that of WT rats. Together, these results suggest that intrinsic changes within the dPAG neurons of *Nlgn3^-/y^* rats underlie the preference for flight responses seen in their behaviour.

In conclusion, we describe altered fear responses in *Nlgn3^-/y^* rats, and provide evidence that this is caused by a circuit bias that predisposes flight over freeze responses. Additionally, we have shown the first phenotypic link between PAG dysfunction and ASD/ID, further study of which may provide additional insight into the mechanisms behind anxiety disorders and abnormal emotional responses seen in people with ASD/ID.

## Methods

### Resource availability

#### Lead contact

Further information and requests for resources and reagents should be directed to and will be fulfilled by the Lead Contact, Prof. Peter Kind.

#### Materials availability

This study did not generate new unique reagents.

### Experimental models and subject details

*Nlgn3*, transgenic rats were bred onto the Sprague-Dawley (SD) background by Horizon Discovery (RRID: RGD_11568700). Rats were housed on either a 14/10 hr (Bangalore Biocluster) or 12/12 hr (University of Edinburgh) light/dark cycle with food and water *ad libitum.* Room temperature was maintained at 21 ± 2°C. Animal husbandry was carried out by University of Edinburgh or Bangalore Biocluster technical staff. Rats were housed 4 rats per cage (2 WT, 2 *Nlgn3^-/y^*, littermates where possible), except for rats that had undergone surgeries for electrode implants, which were single housed in individually ventilated cages (IVCs). All other rats were housed in conventional non-enriched cages. Body weight was monitored throughout experiments.

Rats were handled for a minimum of 3 days prior starting behavioural testing. Each described protocol uses rats that have not experienced any fear-related paradigms before testing, however animals undergoing fear conditioning and active place avoidance tasks underwent several behavioural tasks prior to those shown in this study. These were: marble burying, object recognition memory tasks and three-chamber task.

All procedures were performed in line with the ARRIVE guidelines and both the University of Edinburgh and Home Office guidelines under the 1986 Animals (Scientific Procedures) Act, and CPCSEA (Government of India) and approved by the Animal Ethics Committee of the Institute for Stem Cell Science and Regenerative Medicine (inStem).

Male littermates were assigned to experimental groups based on genotype in order to achieve balanced cohorts. Genotyping was carried out by Transnetyx Inc. at both experimental sites. All experiments and analyses were performed blind to genotype.

### Method details

#### RNA-sequencing

For isolation of RNA, P60-90 male WT and *Nlgn3^-/y^*, rats were anaesthetised with gaseous halothane and decapitated. The brain was quickly extracted and cooled in ice-cold (> 4°C) carbogenated (bubbled with 95% O_2_/ 5% CO_2_) high sucrose artificial cerebrospinal fluid (ACSF) (87 mM NaCl, 2.5mM KCl, 25 mM NaHCO_3_, 1.25 mM NaH_2_PO_4_, 25 mM glucose, 3.4 M sucrose, 7 mM MgCl_2_, 0.5 mM CaCl_2_). The cerebellum was removed, and the brain cut coronally in half before slicing medial-prefrontal cortex. Slices were snap frozen on dry ice and stored at −80°C.

As described in (Hasel *et al.*, 2017), RNA was isolated using Qiagen RNEasy Lipid Tissue kit, with RNA integrity values determined using an Agilent 2100 Bioanalyzer and RNA 6000 Nano chips, with all RIN values 8 or higher. RNA-seq libraries were prepared by Edinburgh Genomics from 1 μg total RNA using the Illumina TruSeq stranded mRNA-seq kit as per the manufacturer’s instructions. Libraries were pooled and sequenced to 50 base paired-end on the Illumina NovaSeq platform to a depth of ~46 million paired-end reads per sample. Reads were mapped to the rat reference genome using STAR RNA-seq aligner version 2.4.oi (Dobin *et al.*, 2013). Read counts per gene were generated from mapped reads with featureCounts version 1.6.3 (Liao *et al.*, 2014), using gene annotations from Ensembl version 82 (Yates *et al.*, 2020).

#### Western Blotting

P60-90 male WT and *Nlgn3^-/y^* rats were anaesthetised with isoflourane and decapitated. The brain was quickly extracted and cooled in ice-cold, carbogenated cACSF. Cortical tissue was dissected, snap frozen on dry ice and weighed. Tissue was then transferred to Bead Mill tubes (Fisher Scientific) containing ice-cold lysis buffer (150 mM NaCl, 1% Triton-X, 0.5% sodium deoxycholate, 0.1% SDS, 50 mM Tris, protease inhibitors, phosphatase inhibitor cocktail sets II and III) (1-5 mg tissue per ml of lysis buffer) and homogenised using a Bead Mill (Fisher Scientific). Laemmli buffer (0.004% bromophenol blue, 10% β-mercaptoethanol, 10% glycerol, 4% SDS, 0.125M Tris-HCl) was added to reach a 1 x concentration with the volume of cell lysate present. Samples were boiled (95°C, 5 minutes), centrifuged (16000 G, 5 minutes) and vortexed.

The protein concentration assay was carried out using a Pierce™ BCA Protein Assay Kit (Fisher Scientific) as per their provided protocol. A logarithmic scale of bovine serum albumin (BSA) standards was made (2 - 0.625 mg/ml). Triplicate samples were incubated at room temperature under agitation. Protein concentrations were then measured using a CLARIOstar plate reader (BMG Labtech), and sample concentrations calculated based on the BSA standard curve.

Samples or protein ladder (PageRuler Plus Prestained Protein Ladder, Fisher Scientific, diluted in Laemmli buffer) was loaded into wells of 10% Mini-PROTEAN TGX Precast Protein Gels (Bio-rad), submersed in running buffer (Bio-rad). The gels were run at a 50 V constant voltage for 30 minutes, then increased to 150 V for 1 hour. The gel was then removed and washed in transfer buffer (Bio-rad) for 15 minutes. Filter paper, nitrocellulose membrane (Bio-rad), sponges and cassettes were washed in water and soaked in transfer buffer prior to transfer. Samples were transferred to the nitrocellulose membrane by running in transfer buffer at 85 V for 2 hours.

The membrane was blocked using blocking buffer (Li-Cor) for 1 hour at room temperature, before incubation with primary antibodies (anti-NLGN3 C-terminus, Synaptic Systems, 1:1000, RRID: AB_2619816.; anti-NLGN3 N-terminus, Novus Biologicals, 1:1000, RRID: AB_11027178) in blocking buffer with 0.01% sodium azide for 10 minutes at room temperature. The membrane was then incubated in secondary antibody (goat anti-rabbit 800, Li-Cor 1:500) in blocking buffer for 2 hours at room temperature. Membrane was then washed in several changes of TBST (TBS: Bio-rad, Tween 20: Sigma Aldrich) for 10 minutes each and subsequently in several changes of TBS, before imaging using an Odyssey infrared imaging system (Li-COR Bioscience).

#### Behavioural paradigms

Rats aged P60-90 were used for all behaviour experiments.

##### Auditory fear conditioning

Fear conditioning (context A) and recall (context B) took place in different contexts in the sound isolation cubicles (Coulbourn Instruments, Whitehall, Pennsylvania, USA). Context A included an aluminium fear conditioning chamber with grid flooring, a black and white horizontal-striped cue. Context B included a cage (35 cm wide, 20 cm deep, 40 cm high) with fresh bedding, mint odour and a transparent Perspex lid. Light conditions were different between contexts; blue-tinted light (~ 5 lux) was used in context A, white/yellow light (~ 20 lux) was used in context B. The behaviour of the animals was recorded using a video camera mounted on the wall of the sound isolation box and a frame grabber (sampling at 30 Hz). Infrared LED cue lights which indicated the onset and termination of auditory cue were placed on the wall of the experimental chambers. A software driven tone generator delivered pure tone using a speaker (4 Ω, Coulbourn Instruments, Whitehall, Pennsylvania, USA) placed on one of the walls inside the experimental chamber. Scrambled foot shock which co-terminated with an auditory cue was delivered by metal grids on the floor connected to shocker box (Habitest system, Coulbourn Instruments, Whitehall, Pennsylvania, USA). The apparatus was cleaned with 70 % ethanol before after the experiments.

Context habituation involved exploration of context B for 20 minutes on 2 consecutive days. On day 3, the rats were subjected to auditory fear conditioning in context A. After a baseline exploration time of 2 minutes in context A, rats were presented with 3 pairings of conditioned stimulus (CS) a continuous tone (CS: 30 s, 5 KHz, 75 dB) which co-terminated a scrambled foot shock as the unconditioned stimulus (US, 0.9 mA for 1 sec). Each CS-US pairing was separated by inter-tone interval (ITI) of 1 minute (modified from Twining *et al.*, 2017).

On days 4 and 5, to determine fear memory recall and extinction, rats were given 2 minutes to explore context B, then presented with 13 CS, with a 30 s ITI. Freezing was scored on all days during all the sessions. Freezing response was evaluated during pre-tone, tone and ITI.

##### Contextual fear conditioning

Rats were introduced to context A and given 2 minutes to explore. They were then presented with 3 unconditioned stimuli (US) pairings (0.9 mA scrambled foot shock for 1 s), with a 1.5 minute ITI. The following day, rats were reintroduced to context A for 10 mins and fear behaviour was scored.

##### Active place avoidance

The rotating platform / carousel maze (Biosignal group, Brooklyn,USA) has a rectangular grid floor (100*100 cm) placed on a circular aluminium base run by an arena motor. This is located 90 cm above ground. This steel grid floor was connected to the constant DC current source box for the shock delivery. This set up was connected to the control box which controls the arena speed via a software interface (Tracker, Biosignal group, Brooklyn, USA). A circular fence made of transparent Perspex was placed over the platform (diameter: 77 cm, height: 32 cm). For data shown in **Figure 3**, a transparent lid was placed on top of the circular fence to prevent the escape of rats. The delivery of foot shocks (0.2 mA, 500 ms, 1500 ms ISI) was tracking based (Carousel Maze Manager; Bahník, 2014). The 60° shock zone was located on either North or South region. The position of the shock zone was counterbalanced between rats, and the rats were placed in the arena facing the wall in one of the adjacent quadrants to the shocked zone. The room consisted of 3 black curtains on three sides and had distinct 3-dimensional cues located at different distances from the apparatus.

Rats were brought to the experimental room and held in a holding cabinet for 30 minutes before the experiment. They were habituated to the rotating arena (as described in Sparks and Reilly, 2016) (1.5 RPM, 2 trials, ITI 10 min in opaque bucket). The following day, rats were given two training sessions over two consecutive days (8 trials per session) in which the 60° shock zone was active. On day 4 a single probe trial was given to animals without shock zone to assess their avoidance memory.

An overhead ceiling camera (Firewire) connected to a framegrabber (DT3155) which recorded and digitised analogue video, feeding it to the tracker software (Biosignal group, Brooklyn, USA). Post-acquisition, the room track and arena track .dat files were analysed in Track Explorer software package (Biosignal group, Brooklyn, USA).

##### Shock-ramp test

Rats were placed within context A from the fear conditioning task. The rats were given 2.5 minutes to explore their environment, then were presented with 3 scrambled foot shocks of 0.06 mA for 1 second per shock, with 1.5 minutes between each shock. This was repeated with the foot shock intensity increased in increments (0.06, 0.1, 0.2, 0.3, 0.5, 0.7, 1 mA). Paw withdrawal, backpedalling, forward or backward running and jumping behaviours were quantified.

### In vivo recording/stimulation of the PAG

#### Implantation of local-field potential electrodes or stimulating electrodes

P60-90 rats were anesthetised with a mixture of isoflurane and O_2_ and their head shaved and sterilised. Each animal was placed on a heat-mat set to 37°C then mounted in a stereotaxic apparatus using atraumatic ear bars. Viscotears™ was applied to the eyes and 4 mg/kg Rimadyl analgesic was injected subcutaneously. Surgery was then performed under aseptic conditions. Paw withdrawal reflexes were checked regularly throughout the surgery and level of isofluorane adjusted accordingly.

A midline scalp incision was made, and craniotomies performed to allow electrode implantation in the PAG (approximate coordinates: bregma −7.46 mm, ventral 4.2 mm, 1 mm lateral from midline).

Recording electrodes (made in-house, ~0.5 mm, 140 μm diameter Teflon coated stainless-steel, A-M systems, USA) or bipolar stimulating electrodes (MS303/3-B/SP, Bilaney Ltd.) were stereotaxically lowered through the craniotomy(ies) to the PAG.

Recording electrodes were implanted unilaterally and affixed to the skull using a combination of UV activated dental cement (SpeedCem, Henry Shein), SuperBond (SunMedical, Japan) and dental cement (Simplex Rapid, Kemdent, UK) then connected to an electronic interface board (EIB 16, Neuralynx). Four miniature screws (Screws and More, Germany) were also attached to the skull for additional support for the recording assembly and to serve as recording ground. Stimulation electrodes were implanted bilaterally and secured to the skull using the same methods as for recording. The incision was closed using absorbable surgical sutures and sterilised with iodine. Rats were left to recover for a minimum of 1 week post-surgery prior to start of the experiment.

#### LFP recordings during fear conditioning

Recordings were made via a 16-channel digitising headstage (C3334, Intan Technologies, USA) connected to a flexible tether cable (12-pin RHD SPI, Intan Technologies, USA), custom built commutator and OpenEphys acquisition board (OEPS, Portugal). LFP signals bandpass-filtered from 0.1-600 Hz and sampled at 2 kHz in OpenEphys software. Rats implanted with LFP electrodes underwent auditory fear conditioning as described above. However, a tone habituation session was carried out in a subset of animals (WT n = 5, KO n = 7) of three 30s tones (5 kHz, 75 dB) with 1 minute ITI, was also added before the conditioning, in order to observe if tone-evoked LFPs were present to an unconditioned tone. Video recordings were made using Freeze Frame software (15 frames per second, Actimetrics) synchronised with electrophysiological signals using TTL pulses.

#### In vivo PAG stimulation

Rats implanted with stimulating electrodes were placed inside context B arena as described for the fear conditioning paradigm. Rats were allowed to explore the arena for 2 minutes, then stimulation (0.1 ms pulses, 100 Hz, 2 seconds) began at an intensity of 30 μA (DS3 isolated constant current stimulators, Digitimer Ltd.) and increased in 5 μA steps up to a maximum of 75 μA (Kim *et al.*, 2013). with an ISI of 3 minutes. Behavioural responses were recorded throughout the protocol using Freeze Frame software.

#### Histology

Following behavioural testing, rats implanted with recording or stimulating electrodes were anesthetised with gaseous isofluorane and intraperitoneal injection of pentobarbitol (27.5 mg/kg) until hindpaw reflexes could not be seen. The electrode sites were lesioned by passing a current pulse of 100 μA for 2 seconds (DS3 isolated constant current stimulators, Digitimer Ltd.) through the headstage. Rats were then transcardially perfused with phosphate-buffered saline, followed by 4% paraformaldehyde (PFA). The brains were extracted and left in 4% PFA for 24 hours. Brains were then cut into 80 μm sections on a vibratome or freezing microtome, and these sections mounted onto glass slides. Sections were then stained with cresyl violet acetate, covered with DPX mounting medium and coverslipped. A Leica DMR upright bright-field microscope was used to image the lesion site. Location of the lesion site was projected onto a schematic of the PAG.

### Ex vivo whole-cell patch clamp recordings

#### Acute slice preparation

Rats aged 4-6 weeks were anaesthetised with gaseous halothane or isofluorane and decapitated. The brain was quickly extracted and cooled in ice-cold (> 4°C) carbogenated (95% O_2_/ 5% CO_2_) high sucrose artificial cerebrospinal fluid (ACSF) (87 mM NaCl, 2.5mM KCl, 25 mM NaHCO_3_, 1.25 mM NaH_2_PO4, 25 mM glucose, 3.4 mM sucrose, 7 mM MgCl_2_, 0.5 mM CaCl_2_). The cerebellum was removed, and the brain cut coronally in half before slicing the PAG coronally at 0.05 mm/s into 400 μm slices on a Leica VT 1200S vibratome. Slices were allowed to recover in carbogenated sucrose-ACSF at 35 ± 1°C for 30 minutes, and then stored at room temperature until recording.

#### Whole-cell recordings

Slices were transferred to a recording chamber where they were perfused with carbogenated recording-ACSF (125 mM NaCl, 2.5 mM KCl, 25 mM NaHCO_3_, 1.25 mM NaH_2_PO_4_, 25 mM glucose, 1 mM MgCl_2_, 2 mM CaCl_2_) at 31 ± 1°C at a rate of 3-6 ml/min. Slices were visualised using infrared differential interference contrast (IR-DIC) video microscopy, using a digital camera (DAGE-MTI) mounted on an upright microscope (U-CA, Olympus, Japan) and a 40x water immersion objective was used for all experiments. These were paired with Scientifica slicescope, patchstar and heater units and controlled using LinLab 2 (Scientifica).

Electrodes with 3-6 MΩ tip resistance were pulled from borosilicate glass capillaries (1.7 mm outer/1mm inner diameter, Harvard Apparatus, UK) horizontal electrode puller (P-97, Sutter Instruments, CA, USA). A potassium-gluconate based internal solution (120 mM K-gluconate, 20 mM KCl, 10 mM HEPES, 4 mM NaCl, 4 mM Mg2ATP, 0.3 mM Na2GTP, pH 7.4, 290-310 mOsm) was used for all current clamp recordings. A caesium-gluconate based internal solution (140 mM Cs-gluconate, 3 mM CsCl, 10 mM HEPES, 0.2 mM EGTA, 5 mM QX-314 chloride, 2 mM MgATP, 0.3 mM Na2GTP, 2 mM NaATP, 10 mM phosphocreatine, pH 7.4, 290-310 mOsm) was used for all voltage-clamp recordings.

Cells in the dorsal and ventral PAG were identified by area. A −70 mV holding potential was applied following the creation of a >1 GΩ seal. The fast and slow membrane capacitances were neutralised before breaking through the cell membrane to achieve whole-cell configuration. For mEPSC recordings, gap-free recordings were performed in voltage-clamp configuration for 10 minutes in the presence of 50 μM picrotoxin and 300 nM tetrodotoxin. Cells were discarded if access resistance was >30 MΩ or changed by >20%. Intrinsic property recordings were carried out in current-clamp configuration, as follows. Resting membrane potential (RMP) of the cell was recorded with current clamped at 0 pA, and all other protocols recorded with appropriate current injection to hold the cell at −70mV. Cells were discarded if RMP was more depolarised than −40 mV or if access resistance was >30 MΩ or changed by >20%. Input resistance and membrane time constant were assessed by injecting a −10 pA step, and cell capacitance calculated from these values. Input-output curves and rheobase potential was assessed by current injections of −200 to +100 pA for 500 ms (10 pA steps). Action potential kinetics were gleaned from the rheobase action potential. Recordings were attained using a Multiclamp 700B amplifier linked to pCLAMP™ Clampex software (Molecular Devices). Signals were sampled at 20 kHz (Digidata1440 or Digidata1550A, Molecular Devices) and Bessel-filtered at 2 kHz for voltage-clamp recordings and 10 kHz for current-clamp recordings.

### Quantification and statistical analysis

Fear behaviour was scored as either ‘classical freezing’, defined as no movement except for respiration (Blanchard and Blanchard, 1989), or ‘paw immobility response’, defined as all 4 paws unmoving, however allowing for movement of the head and neck. Each of these behaviours was scored if lasting > 1 second. For the shock sensitivity paradigm, occurrence of paw withdrawal responses, back-pedalling, forward/backward running and jumping were scored. For dPAG stimulation experiments, response behaviours were scored as freezing, startle, attention, running, or jumping, according to criteria described by Calvo *et al.* (2019). All behaviour was manually scored using BORIS software (Friard and Gamba, 2016) or in-house software Z-score (created by O. Hardt).

Stimfit software (Guzman *et al.*, 2014) plus custom-written Matlab scripts (A. Jackson) were used for whole-cell patch-clamp data analysis. mEPSCs were analysed for the final 3 minutes of the 10 minute recording. Events were detected using template-matching and filtered to at 3 x standard deviation of baseline (Clements and Bekkers, 1997).

Data collected from LFP recordings were analysed using custom-written Matlab scripts (F. Inkpen, A. Jackson). Raw traces of 3 tones were averaged, and then z-scored to normalise data to baseline noise. Peak and trough of the LFPs were manually selected.

Throughout, all data is shown as mean ± SEM, or as percentages where appropriate. Statistics were carried out using GraphPad Prism software 8.0, SPSS, or RStudio. Two-way ANOVAs with Holm-Sidak post-hoc repeated measures test (Figures 2, 3, 4, 5, 6, 7, Supplemental Figure 1, 2, 4), unpaired t-tests (Figure 4), paired t-tests (Supplemental Figure 3), Fisher’s exact tests (Figures 3, 7), three-way ANOVAs (Figures 2, 6, Supplemental Figure 1), or generalised linear mixed modelling (GLMM) (Figure 5, Supplemental Figure 4) were employed. N was taken to be animal average in all cases to avoid pseudoreplication, except for when GLMM statistical analysis was employed. R packages lme4 and car were utilised to perform GLMMs. P values of <0.05 were taken to be significant, and one star (*) represents all p values <0.05 throughout.

## Author contributions

Conceptualisation, PCK, OH, ERW, NJA, TCW, VK, and ST; Methodology, TCW, AB, DJAW, SC, ERW, OH, and PCK; Software, OH, FHI, and ADJ; Formal analysis, ORD, ZK, XH, NJA, VK, and ST; Investigation, NJA, VKm ST, TCW, AKHT PB, MSN, and AK; Visualisation, NJA, VK, and ST; Writing - original draft, NJA, and PCK; Writing - review and editing, NJA, VK, TCW, ORD, AB, DJAW, SC, ERW, OH, and PCK; Funding acquisition, AB, DJAW, SC, ERW, OH, and PCK; Supervision, AB, DJAW, SC, ERW, OH, and PCK.

## Acknowledgements

Many thanks to Siddhartha Datta, Suryanarayan Biswal and Urvashi Bhattacharyya for their help with behaviour analysis, and to Lynsey Dunsmore, Arpita Sharma and Priyangvada Singha for colony management and interpreting genotyping results.

