## Supplemental Figures for "Imbalance of flight-freeze responses and their cellular correlates in the *Nlgn3^-/y^* rat model of autism"

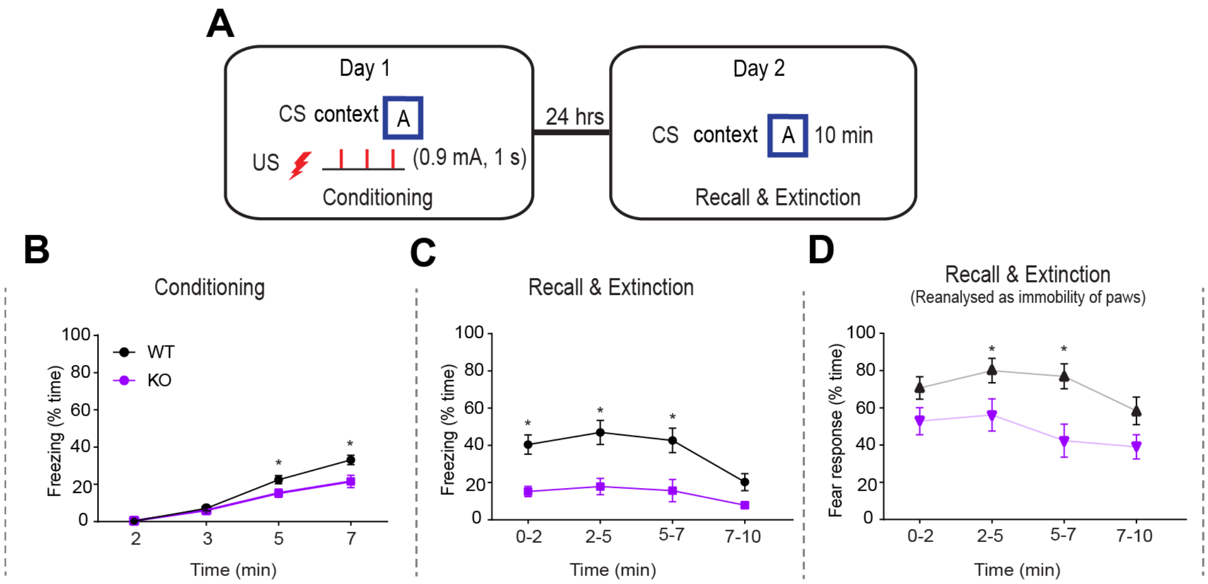


**Supplemental figure 1. *Nlgn3*$^{-/y}$ rats display reduced classic freezing behaviour in a contextual fear conditioning paradigm.** (A) Schematic of contextual fear conditioning paradigm. (B) Classic freezing behaviour is reduced in *Nlgn3*$^{-/y}$ rats in comparison to WTs during the conditioning phase of contextual fear conditioning ($F_{(1, 25)} = 5.67$, p = 0.025, repeated measures two-way ANOVA, WT n = 13, KO n = 14). (C) Classic freezing behaviour is reduced in *Nlgn3*$^{-/y}$ rats in comparison to WTs during the recall phase of contextual fear conditioning ($F_{(1, 25)} = 26.61$, p > 0.0001, repeated measures two-way ANOVA, WT n = 13, KO n = 14). (D) When analysed as "immobility response" (i.e all four paws unmoving but allowing for movement of head and neck, shown in light purple/grey) *Nlgn3*$^{-/y}$ rats show a response to the CS significantly different to classic freezing (main effects of scoring method: p <0.0001, $F_{(1, 25)} = 200.82$, and genotype: p <0.0001, $F_{(1, 25)} = 20.65$, three-way ANOVA, WT n = 13, KO n = 14).

Data represented as mean ± SEM.

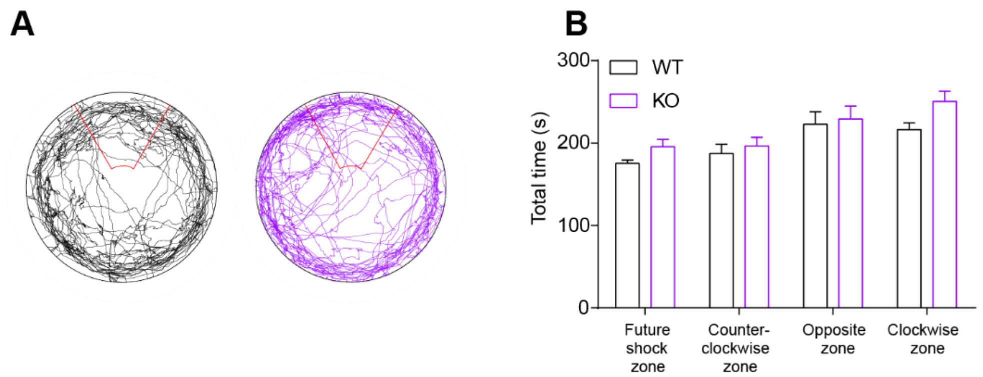


**Supplemental figure 2. *WT and Nlgn3$^{-/y}$* rats explore the APA arena equally during habituation sessions.** (A) Representative track plots from WT (black) and *Nlgn3$^{-/y}$* (purple) rats during habituation to the arena. (B) Total number of seconds WT and *Nlgn3$^{-/y}$* rats spend in each quadrant of the arena over two habituation days (p = 0.069, $F_{(1, 21)}$ = 3.67, repeated measures two-way ANOVA, WT n = 12, KO n = 11).

Data represented as mean ± SEM.

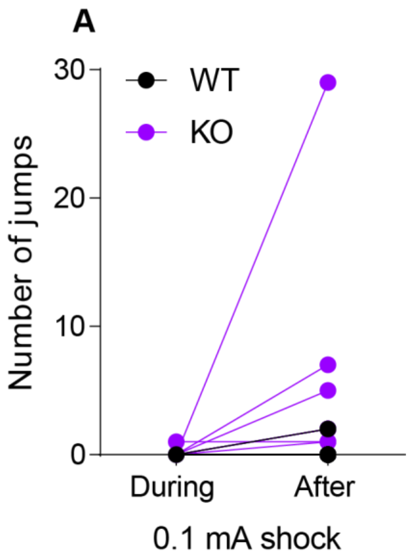


**Supplemental figure 3. Effect of repeated footshocks on WT and *Nlgn3$^{-/y}$* rats.** (A) Number of jumps exhibited in response to 0.1 mA foot shocks during (following 0.06 mA) and after (following 1 mA) shock ramp testing. Number of jumps are not significantly different for WT ($p = 0.35$, paired t-test, $n = 11$) or KO ($p = 0.10$, paired t-test, $n = 14$) animals.

Dots represent individual animals.

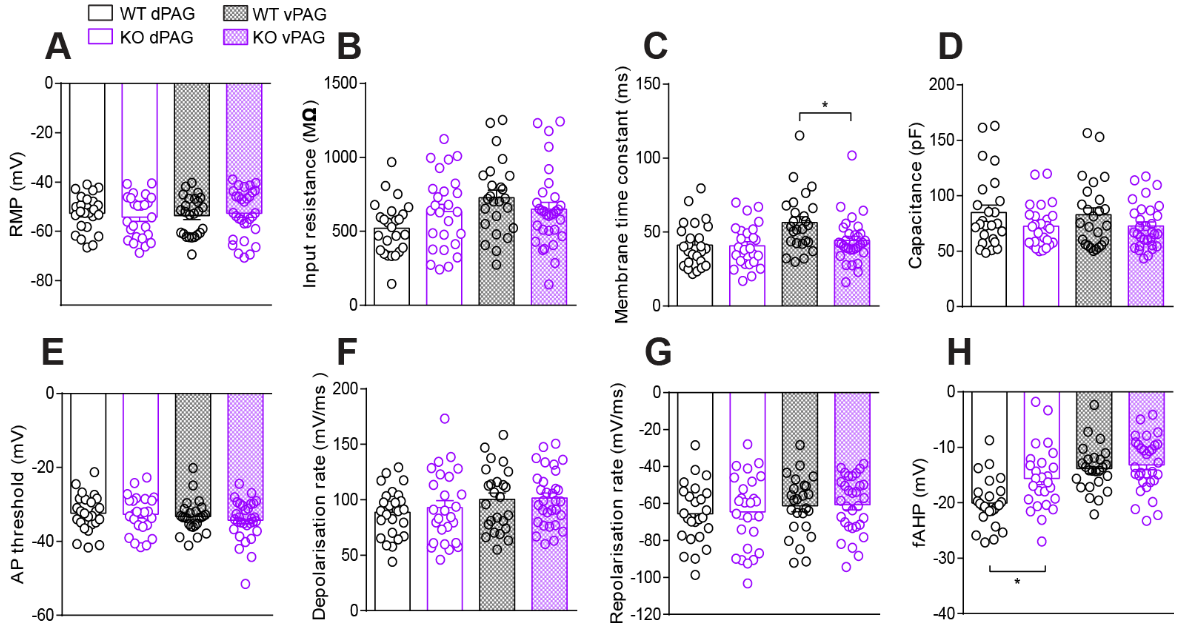


**Supplemental figure 4. Intrinsic properties of PAG cells recorded from WT and *Nlgn3*$^{-/y}$ rats.** (A) Resting membrane potential is comparable between *Nlgn3*$^{-/y}$ and WT rats in both dPAG (p = 0.61, GLMM, dPAG WT, 25 cells/ 10 rats, dPAG KO 26 cells/ 9 rats) and vPAG cells (p = 0.75, GLMM, WT 24 cells/10 rats, vPAG KO 28 cells/ 9 rats). (B) Input resistance is comparable between *Nlgn3*$^{-/y}$ and WT rats in both dPAG  (p = 0.090, GLMM, dPAG WT, 25 cells/ 10 rats, dPAG KO 26 cells/ 9 rats) and vPAG cells(p = 0.26, GLMM, vPAG WT 24 cells/ 9 rats, vPAG KO 28 cells/ 10 rats). (C) Membrane time constant is comparable between *Nlgn3*$^{-/y}$ and WT rats in cells recorded from dPAG (p = 0.78, GLMM, dPAG WT, 25 cells/ 10 rats, dPAG KO 26 cells/ 9 rats), however is reduced in vPAG cells of *Nlgn3*$^{-/y}$ compared to WT (p = 0.0095, GLMM, vPAG WT 24 cells/ 9 rats, vPAG KO 28 cells/ 10 rats). (D) Capacitance is comparable between *Nlgn3*$^{-/y}$ and WT rats in both dPAG (p = 0.11, GLMM, dPAG WT, 25 cells/ 10 rats, dPAG KO 26 cells/ 9 rats) and vPAG cells (p = 0.19, GLMM, vPAG WT 24 cells/ 9 rats, vPAG KO 28 cells/ 10 rats). (E) Action potential (AP) threshold is comparable between *Nlgn3*$^{-/y}$ and WT rats in both dPAG (p = 0.86, GLMM, dPAG WT, 25 cells/ 10 rats, dPAG KO 26 cells/ 9 rats) and vPAG cells (p = 0.47, GLMM, vPAG WT 24 cells/ 9 rats, vPAG KO 28 cells/ 10 rats). (F) No difference in AP depolarisation rate between WT and *Nlgn3*$^{-/y}$ rats in either dPAG (p = 0.71, GLMM, dPAG WT, 25 cells/ 10 rats, dPAG KO 26 cells/ 9 rats) or vPAG cells (p = 0.90, GLMM, vPAG WT 24 cells/ 9 rats, vPAG KO 28 cells/ 10 rats). (G) No difference in AP repolarisation rate between WT and *Nlgn3*$^{-/y}$ rats in either dPAG (p = 0.76, GLMM, dPAG WT, 25 cells/ 10 rats, dPAG KO 26 cells/ 9 rats) or vPAG cells (p = 0.90, GLMM, vPAG WT 24 cells/ 9 rats, vPAG KO 28 cells/ 10 rats). (H) Fast afterhyperpolarisation potential (fAHP) is significantly reduced in *Nlgn3*$^{-/y}$ rat dPAG neurons in comparison to WT (p = 0.0047, GLMM, dPAG WT, 25 cells/ 10 rats, dPAG KO 26 cells/ 9 rats) but unchanged in vPAG neurons (p = 0.58, GLMM, vPAG WT 24 cells/ 9 rats, vPAG KO 28 cells/ 10 rats).

Data represented as mean ± SEM, dots represent individual cells.

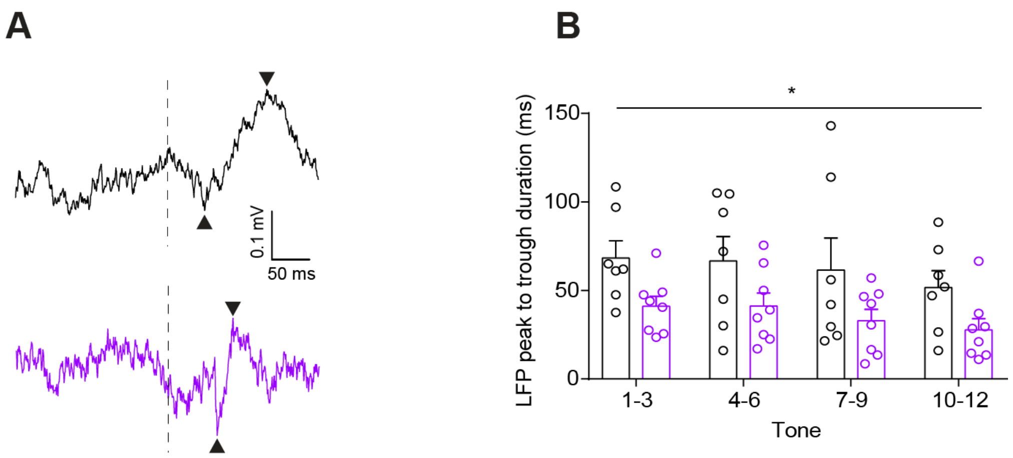


**Supplemental figure 5. PAG ERPs during fear recall are significantly shorter duration in *Nlgn3$^{-/y}$* rats.**

(A) Example LFP traces from WT (black) and *Nlgn3$^{-/y}$* (purple) rats. Black arrows denote trough and peak. (B) *Nlgn3$^{-/y}$* rats display significantly faster CS-evoked ERPs in the PAG during fear recall in comparison to WT rats ($p = 0.042$, $F_{(1, 13)} = 5.09$, two-way ANOVA, WT n = 7, KO n = 8).

Data represented as mean ± SEM, dots represent individual animals.

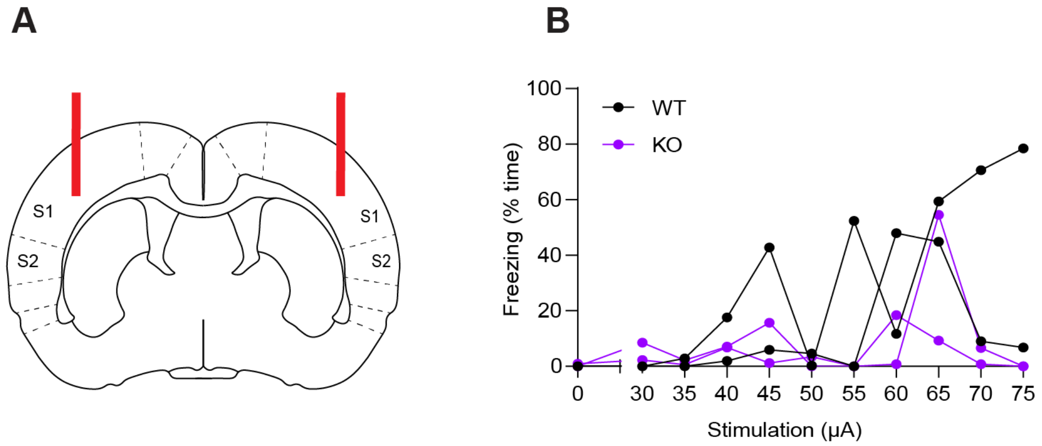


**Supplemental figure 6. Defensive reactions were not elicited by electrical stimulation of primary somatosensory cortex in WT or *Nlgn3*$^{-/y}$ rats.** (A) Schematic depicting stimulating electrode (red lines) implant site. (B) Freezing behaviour, defined as no movement except for respiration, for 2 WT and 2 *Nlgn3*$^{-/y}$ rats receiving cortical stimulation. Resting or sleeping was indistinguishable from freezing given this definition.

Connected lines represent an individual animal, points represent average freezing time for 3 minutes post-stimulation.
